# Activation of Sarm1 produces cADPR to increase intra-axonal calcium and promote axon degeneration in CIPN

**DOI:** 10.1101/2021.04.15.440024

**Authors:** Yihang Li, Maria F. Pazyra-Murphy, Daina Avizonis, Mariana de Sa Tavares Russo, Sophia Tang, Johann S. Bergholz, Tao Jiang, Jean J. Zhao, Jian Zhu, Kwang Woo Ko, Jeffrey Milbrandt, Aaron DiAntonio, Rosalind A. Segal

## Abstract

Cancer patients frequently develop chemotherapy-induced peripheral neuropathy (CIPN), a painful and long-lasting disorder with profound somatosensory deficits. There are no effective therapies to prevent or treat this disorder. Pathologically, CIPN is characterized by a “dying-back” axonopathy that begins at intra-epidermal nerve terminals of sensory neurons and progresses in a retrograde fashion. Calcium dysregulation constitutes a critical event in CIPN, but it is not known how chemotherapies such as paclitaxel alter intra-axonal calcium and cause degeneration. Here, we demonstrate that paclitaxel triggers Sarm1-dependent cADPR production in distal axons, promoting intra-axonal calcium flux from both intracellular and extracellular calcium stores. Genetic or pharmacologic antagonists of cADPR signaling prevent paclitaxel-induced axon degeneration and allodynia symptoms, without mitigating the anti-neoplastic efficacy of paclitaxel. Our data demonstrate that cADPR is a calcium modulating factor that promotes paclitaxel-induced axon degeneration and suggest that targeting cADPR signaling provides a potential therapeutic approach for treating CIPN.

**HIGHLIGHTS:**

- Paclitaxel induces intra-axonal calcium flux
- Sarm1-dependent cADPR production promotes axonal calcium elevation and degeneration
- Antagonizing cADPR signaling pathway protects against paclitaxel-induced peripheral neuropathy *in vitro* and *in vivo*

## INTRODUCTION

Axons provide critical long-distance connections among neurons. Functional and structural defects in these connections, known as axonopathy, are a hallmark of many central and peripheral neurodegenerative diseases. Toxins, disease or injury can all initiate local signaling cascades leading to dysfunction and destruction of the affected axon. Often axon degeneration begins at axon terminals and progresses in a retrograde direction towards the neuron cell bodies, a process known as “dying-back” degeneration (Wang et al., 2012). “Dying-back” axon degeneration is a key pathological feature observed in cancer patients suffering from chemotherapy-induced peripheral neuropathy (CIPN) (Fukuda et al., 2017). The most common manifestations of CIPN are excess pain and deficits of sensation in a “stocking and glove” distribution, in which the distal limbs are most significantly affected (Schneider et al., 2015; Seretny et al., 2014). Although CIPN is often reversible over time, a significant fraction of patients develop long-lasting, untreatable pain and severe sensory deficits persisting for years even after the cessation of cancer treatment (Dorsey et al., 2019). Defective axonal transport, mitochondrial dysfunction and intra-axonal calcium dysregulation have all been implicated as key events in the process of chemotherapy-induced axon degeneration (Berbusse et al., 2016; Bobylev et al., 2015; Boehmerle et al., 2007; Pease-Raissi et al., 2017). However, it remains unclear how these processes coordinate with one another to cause degeneration. Given the lack of understanding of the underlying mechanisms, it is not surprising that there are no effective treatments available for CIPN. Methods of mitigating CIPN are currently limited to dose reduction or early termination of chemotherapy, which compromises the therapeutic response and patient survival (Dorsey et al., 2019). Thus, there is an urgent need for improved understanding of CIPN mechanisms and discovery of effective therapeutic targets.

Recent insights into the mechanism of axon degeneration caused by physical injury provide important clues for understanding axon degeneration due to chemotherapy. Studies of Wallerian degeneration in *Drosophila*, Zebrafish and mammalian models have identified Sterile alpha and TIR motif-containing 1 (Sarm1) as a key mediator of injury-induced axonal degeneration (Gerdts et al., 2015; Gerdts et al., 2013; Osterloh et al., 2012; Tian et al., 2020). Loss of Sarm1 also protects against both diabetes and chemotherapy-induced neuropathy (Geisler et al., 2019a; Geisler et al., 2016; Turkiew et al., 2017), suggesting a convergent pathway among axon degeneration induced by injury, chemotherapy and metabolic disorder. Sarm1 functions as an NADase, hydrolyzing NAD to generate nicotinamide, cyclic-ADP Ribose (cADPR) and ADP Ribose (ADPR) (Essuman et al., 2017). The catalytic domain of Sarm1 is located in the C-terminal TIR domain (Essuman et al., 2017). Activation of the TIR domain by dimerization is sufficient to trigger axon degeneration (Gerdts et al., 2015). Conversely, mutations that enzymatically disable Sarm1 prevent axotomy-induced axon degeneration (Essuman et al., 2017; Geisler et al., 2019b; Summers et al., 2016). These findings indicate the NADase activity of Sarm1 is both necessary and sufficient for its pro-degenerative function. The question remains whether Sarm1-dependent axon degeneration is due to NAD loss and subsequent metabolic catastrophe; or whether NAD-derived metabolites such as cADPR and ADPR also contribute to degeneration. A recent study showed that NAD breakdown products signal degeneration following activation of plant TIR domains (Wan et al., 2019), suggesting excess cADPR and ADPR production can contribute to a degenerative process. cADPR is a calcium mobilizing agent that potentiates calcium-induced calcium release in sea urchin egg homogenate (Clapper et al., 1987; Lee, 1993; Lee et al., 1989), and in other cell types, including T-lymphocytes (Guse et al., 1999), cardiac myocytes (Rakovic et al., 1996), and neurons (Currie et al., 1992; Higashida et al., 2001). Studies using pharmacological approaches or *in vitro* reconstitution of channels have shown that cADPR modulates intracellular calcium release through ryanodine receptor channels (RyRs) but not inositol 1,4,5-trisphosphate receptors (IP_3_Rs) (Clapper et al., 1987; Dargie et al., 1990; Meszaros et al., 1993). Moreover, both cADPR and ADPR activate TRPM2, a calcium permeable non-specific cation channel located predominantly on the plasma membrane (Kolisek et al., 2005; Yu et al., 2019). These observations raise the possibility that the products of activated Sarm1 may contribute to axonal calcium dysregulation.

Axotomy rapidly induces increased axonal calcium flux (Adalbert et al., 2012; Loreto et al., 2015; Vargas et al., 2015); and both extracellular and intracellular calcium stores are involved (Villegas et al., 2014). Paclitaxel and vincristine, two microtubule-targeting chemo-drugs, alter ATP-induced calcium release in neuroblastoma cells and DRG primary cultures (Benbow et al., 2012; Boehmerle et al., 2007; Pease-Raissi et al., 2017), while depletion of calcium channels or inhibition of calpain, a calcium-dependent protease, prevent paclitaxel-induced axon degeneration (Pease-Raissi et al., 2017; Wang et al., 2004). These data collectively identify calcium dysregulation as a key component of axon degeneration induced by both chemotherapy and injury. However, current studies lack direct observation of calcium changes within axons following chemotherapy drug treatment, and have not clearly identified the sources of calcium that contribute to degeneration.

Here we directly visualized intra-axonal calcium changes following paclitaxel treatment using GCaMP6s targeted to axons of sensory neurons in microfluidic chambers. We found that paclitaxel leads to Sarm1 activation and excess cADPR production, which contributes to paclitaxel-induced intra-axonal calcium flux and degeneration. We found that interference with cADPR signaling using genetic or pharmacologic intervention prevents paclitaxel- or Sarm1 activation-triggered axonal calcium elevation and degeneration in cultured sensory neurons. Finally, we showed that a cADPR antagonist attenuates paclitaxel-induced neuropathy *in vivo* without affecting the anti-tumor efficacy of paclitaxel. Our data provide new insights into the molecular mechanisms of paclitaxel-induced peripheral neuropathy and suggest cADPR as a promising therapeutic target for treating CIPN.

## RESULTS

### Paclitaxel treatment increases cADPR levels through Sarm1 activation

Local paclitaxel treatment of the axons of sensory neurons in compartmentalized cultures leads to axon degeneration in DRG neurons (Figure1A) (Gornstein and Schwarz, 2017; Pease-Raissi et al., 2017; Yang et al., 2009). This axon degeneration was quantified as the ratio of fragmented axons normalized to total axon area (Figure1B). In contrast, paclitaxel-induced axon degeneration was prevented in Sarm1 shRNA-transduced DRG neurons (Figure1A-B, FigureS1C). The efficiency of Sarm1 shRNA was verified using RT-qPCR (FigureS1A). This result demonstrates that Sarm1 is required for paclitaxel-induced axon degeneration in a cell autonomous fashion; moreover, the compartmentalized culture system provides a good *in vitro* model to address the mechanisms of Sarm1-dependent axon degeneration.

**Figure1.**
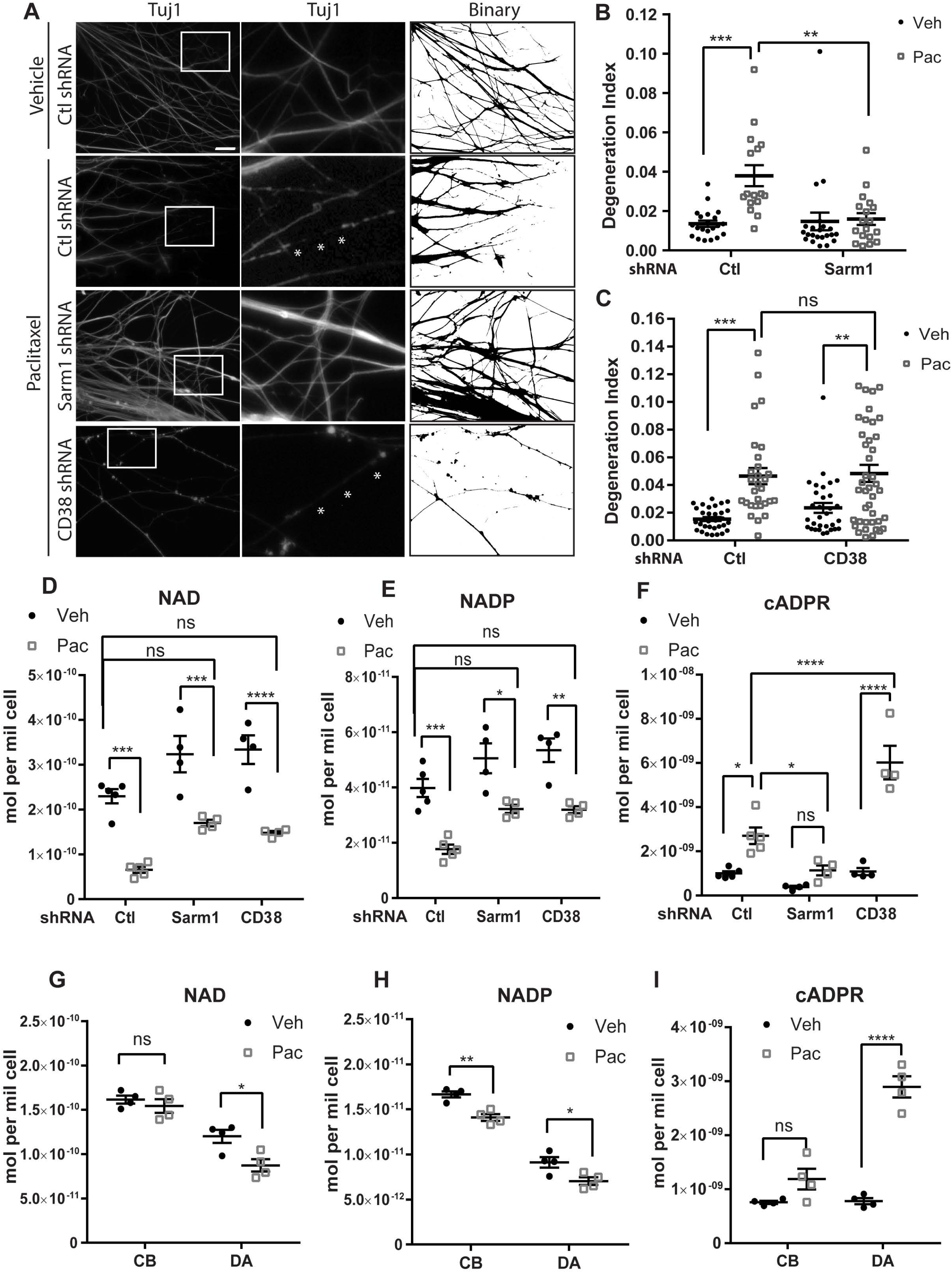
**Paclitaxel-induced cADPR production and axon degeneration require Sarm1 but not CD38.** (A) Tuj1 immunostaining and corresponding binarized images of axon endings of DRG neurons grown in compartmented cultures. Following lentiviral infection with shRNA against Sarm1 or CD38, or control shRNA (Ctl), 30nM paclitaxel (Pac) or vehicle control (Veh) were applied to axons for 24 hours. White boxes outline regions shown at higher magnification in the center panels. White stars indicate fragmented region of axons displayed as interruptions in Tuj1 continuity. Scale bar: 20µm. See FigureS1C for representative images of vehicle treated axons expressing Sarm1 or CD38 shRNA. (B-C) Quantification of the degeneration index of (A) in DRG neurons with Sarm1 knockdown (B) and CD38 knockdown (C): ratio of area of fragmented axons to total axon area (degeneration index). Data represent mean +/- SEM; individual data points are shown. N represents number of images. Data is pooled from three (B) and five (C) independent experiments. *p<0.05, **p<0.01, ***p<0.001, ****p<0.0001 by two-way ANOVA with Tukey’s multiple comparison test. (D-F) Level of NAD (D), NADP (E) and cADPR (F) in DRG neurons cultured for 24hrs with 600nM paclitaxel (Pac) or DMSO (Veh) following lentiviral infection with shRNA to Sarm1 or CD38, or control shRNA. Data represent mean +/- SEM; individual data points are shown. N represents number of independent experiments. *p<0.05, **p<0.01, ***p<0.001, ****p<0.0001 by two-way ANOVA with Tukey’s multiple comparison test. (G-I) Levels of NAD (G), NADP (H) and cADPR (I) in distal axons (DA) and cell bodies (CB) of DRG neurons after 24hrs of 30nM paclitaxel (Pac) or DMSO (Veh) applied to axons. Data represent mean +/- SEM, individual data points are shown. N represents number of independent experiments, *p<0.05, **p<0.01, ****p<0.0001 two-way ANOVA, Tukey’s multiple comparison test.

As activated Sarm1 catalyzes the breakdown of NAD and NADP to generate cADPR and ADPR, we asked whether paclitaxel affects NAD and its metabolites. Paclitaxel treatment of DRG sensory neuron cultures significantly decreased NAD and NADP levels as measured via LC-MS/MS (Figure1D-E). Knockdown of Sarm1 increased NAD and NADP levels in both vehicle and paclitaxel treated cultures, although this did not completely rescue the NAD or NADP depletion caused by paclitaxel (Figure1D-E). Paclitaxel treatment also significantly increased cADPR levels (Figure1F), the product of NAD hydrolysis and a validated biomarker of SARM1 activity (Sasaki et al., 2020). Sarm1 depletion inhibited paclitaxel-induced cADPR production and restored cADPR to baseline levels (Figure1F).

We then asked how these metabolic changes contribute to paclitaxel-induced axon degeneration. One possibility is that excess NAD/NADP consumption leads to insufficient energy supply and axon maintenance failure, thereby triggering degeneration. To investigate whether NAD/NADP levels are critical for paclitaxel-induced axon degeneration, we tested the neuroprotective effect of CD38, another NADase (Aksoy et al., 2006). Knockdown of CD38 using shRNA restored NAD and NADP to the levels seen in control cultures (Figure1D-E, FigureS1B). However, CD38 depletion failed to block paclitaxel-induced degeneration (Figure1A,C, FigureS1C). This result is consistent with the previous finding that murine *CD38-/-* neurons were not protected from axotomy (Sasaki et al., 2009). These data suggest that NAD/NADP levels are not critical determinants of paclitaxel-induced axon degeneration.

Surprisingly, rather than inhibiting cADPR production, CD38-deficient DRG neurons showed enhanced cADPR elevation when treated with paclitaxel (Figure1F). This was a synergistic effect with paclitaxel, as CD38 depletion alone did not alter basal levels of cADPR (Figure1F). Since CD38 functions as both an NADase and a cADPR hydrolase (Chini, 2009), it is possible that excess cADPR produced by paclitaxel-induced Sarm1 activity accumulates further upon CD38 deficiency. The differential effects of Sarm1 and CD38 on cADPR production raises the possibility that cADPR itself contributes to paclitaxel-induced degeneration.

We then evaluated the subcellular localization of paclitaxel-induced metabolites using sensory neurons in the compartmentalized culture systems. Axonal treatment with paclitaxel significantly increased cADPR levels in distal axons but not in cell bodies (Figure1I), while NAD and NADP levels showed small decreases in both distal axons and cell bodies (Figure1G-H). We also found that ADPR, the other breakdown component of NAD, was not affected by paclitaxel in either cell bodies or distal axons (FigureS1D). These results demonstrate that the local accumulation of cADPR in axons correlates with paclitaxel-induced axon degeneration.

### Paclitaxel leads to Sarm1-dependent axonal calcium flux

Calcium dysregulation is a critical step in the process of axon degeneration. However, there is no direct evidence showing paclitaxel changes axonal calcium flux. As cADPR can potentiate calcium release from intracellular stores (Clapper et al., 1987; Lee et al., 1989), the paclitaxel-induced increase in cADPR in distal axons could lead to changes in intra-axonal calcium flux. To directly monitor axonal calcium alterations, we cultured DRG neurons in microfluidics and transduced the neurons with AAV9 expressing axon localizing GCaMP6s co-translated with mRuby3 (AAV9-GCaMP6s-mRuby3) (Broussard et al., 2018), together with lentivirus expressing Sarm1 shRNA or tGFP targeted control shRNA. Of note, the substrate in microfluidic cultures consists of laminin, rather than the Matrigel used in Campenot cultures. A previous study suggests that changes in substrate may alter the timing and dose responses of paclitaxel induced axon degeneration (Shin et al., 2021). We used a higher dose of paclitaxel which is required in non-compartmental cultures (600nM) and also causes axon retraction in microfluidic chambers (FigueS2A-B). We imaged GCaMP6s and mRuby3 signals over the course of 48 hours of paclitaxel or vehicle treatment. At each time point the GCaMP6s signal was normalized to mRuby3 signal (GCaMP6s/mRuby3) and this ratio was then normalized to GCaMP6s/mRuby3 at time 0 [(GCaMP6s/mRuby3)0]. We found that paclitaxel gradually increases axonal calcium signal (Figure2A-B), which is followed by axon degeneration (FigureS2C, white arrows).

**Figure2.**
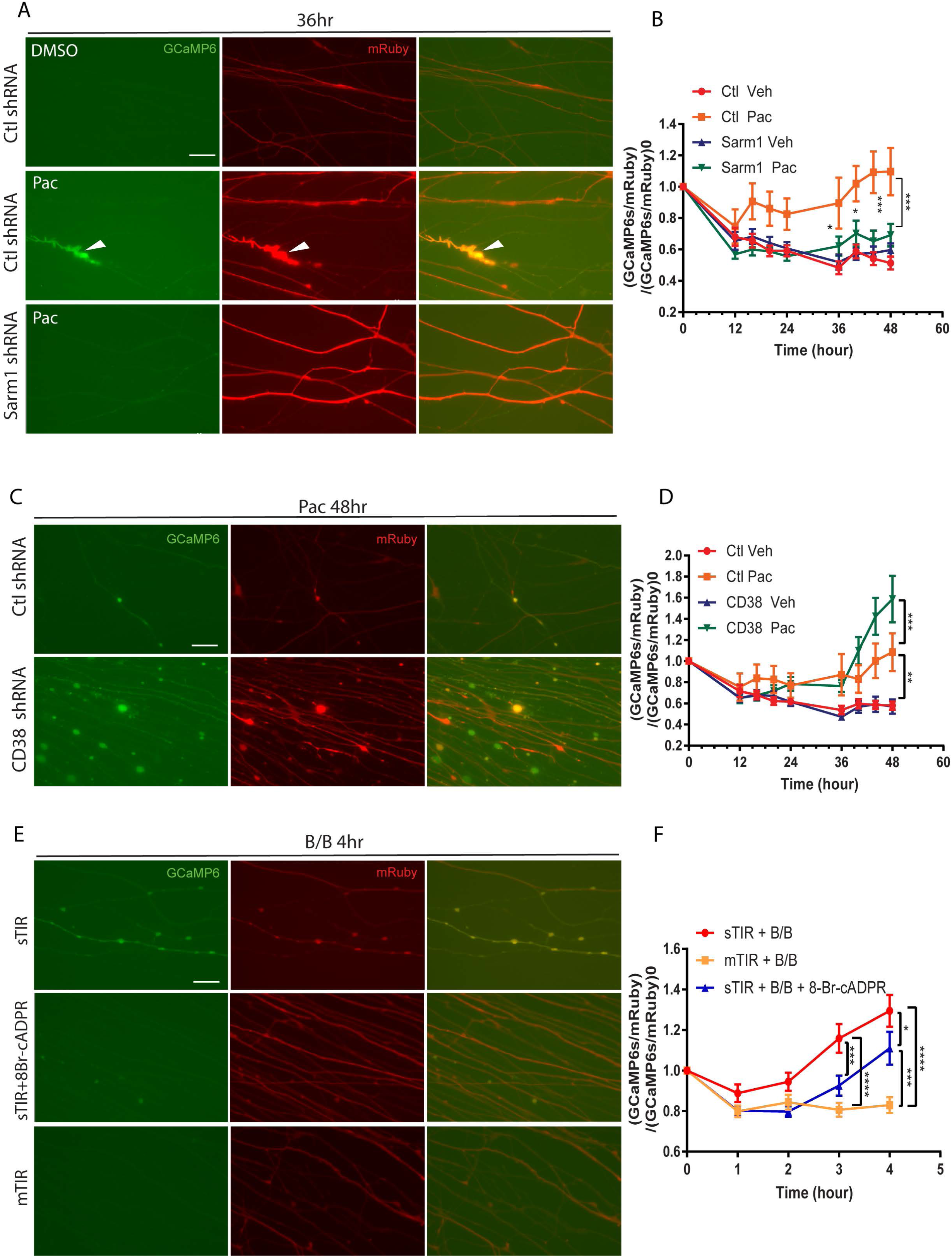
**Paclitaxel leads to Sarm1-dependent increase in axonal calcium.** (A, C) Representative images of GCaMP6s (green) and mRuby3 (red) in axons of DRG neurons also expressing indicated shRNA at 36hrs (A) and 48hrs (C) after 600nM paclitaxel (Pac) or vehicle control (Veh) was applied to axons. White arrow indicates axon with elevated calcium signal. This axon later degenerates (also see FigureS2A). Scale bar: 20µM. (B, D) Quantification of calcium signal calculated as (GCaMP6s/mRuby3)/(GCaMP6s/mRuby3)_0_. (GCaMP6s/mRuby3) represents calcium signal at each time point and (GCaMP6s/mRuby3)_0_ represents calcium signal at time 0 before treatment. Data represent mean +/- SEM. N represents number of images. 8-10 images from each sample per experiment; 1-2 samples per condition per experiment. Data pooled from four independent experiments. Three experiments from (D) share the same control shRNA samples with (B). *p<0.05, **p<0.01, ***p<0.001, ****p < 0.0001 by two-way ANOVA with Tukey’s multiple comparison test for each time point. (E) Representative images of GCaMP6s (green) and mRuby3 (red) of axons of DRG neurons expressing FkbpF36V-sTIR (top and middle) or FkbpF36V-mTIR (bottom) 4hrs after axonal treatment of 50nM B/B homodimerizer together with 10µM 8-Br-cADPR (middle) or saline (top and bottom). Scale bar: 20µM (F) Quantification of calcium signal of (E). Data represent mean +/- SEM. N represents number of images. 8 images from each sample per experiment; 2 samples per condition per experiment. Data pooled from four independent experiments. *p<0.05, ***p<0.001, ****p < 0.0001 by two-way ANOVA with Tukey’s multiple comparison test for each time point.

In sensory neurons, knockdown of Sarm1 prevented paclitaxel-induced axonal calcium elevation (Figure2A-B and FigureS2C), indicating that Sarm1 is required for paclitaxel-induced axonal calcium flux. As CD38 depletion enhances the increase in cADPR levels induced by paclitaxel, we also examined the calcium signal in CD38-depleted cultures. In direct contrast to Sarm1 knockdown, CD38 knockdown actually increased axonal calcium levels in axons treated with paclitaxel, but did not alter calcium signals in axons treated with vehicle control (Figure2C-D). These results indicate that changes in cADPR correlate well with axonal calcium levels.

To determine if Sarm1 activation is sufficient to increase axonal calcium flux, we transduced DRG neurons with AAV9-GCaMP6s-mRuby3 together with lentivirus expressing FkbyF36V tagged Sarm1 TIR domain (sTIR). Dimerization of this TIR domain activates the enzyme, generating cADPR (FigureS2D), and causes axon degeneration within 2-6 hours (Gerdts et al., 2015). Axonal calcium signals started to increase approximately 2 hours after B/B homodimerizer was added to the axon chamber and signal continued to increase throughout the 4 hour time window (Figure2E-F). As a control, dimerization of FkbpF36V tagged MYD88 TIR domain (mTIR), which does not possess NADase activity or induce axon degeneration (Gerdts et al., 2015), failed to trigger intra-axonal calcium elevation (Figure2E-F). These data together indicate that Sarm1 is necessary and sufficient for axonal calcium elevation.

We then asked whether cADPR contributes to paclitaxel induced axonal calcium flux. 8-Br-cADPR is a cell permeable antagonist of cADPR, which blocks cADPR-induced calcium release in sea urchin egg homogenate (Walseth and Lee, 1993) as well as in astrocytes and bone marrow neutrophils (Banerjee et al., 2008; Partida-Sanchez et al., 2007). We found that 8-Br-cADPR treatment in axons partially inhibited axonal calcium elevation caused by sTIR dimerization (Figure2E-F). We tested a second cADPR antagonist 8-Br-7-CH-cADPR, which showed similar inhibition of calcium elevation caused by sTIR dimerization (FigureS2E). Moreover, treatment with 8-Br-cADPR together with paclitaxel in axons also partially decreased the axonal calcium flux compared to axons treated with paclitaxel alone (FigureS2F-G). The partial rescue by 8-Br-cADPR indicates that cADPR contributes to calcium modulation in axons, but it is not the only factor that is involved in paclitaxel-triggered calcium flux. cADPR is known to modulate calcium release in a calcium-induced calcium release manner (Galione et al., 1991). Our data is consistent with a model in which cADPR functions as a calcium modulator that enhances calcium release and synergizes with other factors (Galione and White, 1994) to induce axon degeneration.

### Inhibitors of cADPR dependent calcium signaling pathway is neuroprotective *in vitro*

To investigate this model and determine whether cADPR-dependent calcium mobilization contributes to axon degeneration downstream of paclitaxel, we tested the neuroprotective effect of 8-Br-cADPR in compartmentalized cultures. Axonal treatment with 8-Br-cADPR significantly decreased paclitaxel-induced degeneration in a concentration dependent manner (Figure3A-B). We found that 8-Br-cADPR also protects against axon degeneration induced by activated Sarm1 using sTIR dimerization to activate Sarm1 enzymatic activity. This direct activation of Sarm1 by dimerization with B/B homodimerizer triggered acute axon degeneration in DRG sensory neurons within 3 hours, and this degeneration was inhibited by 8-Br-cADPR (FigureS3A-B). These findings suggest that 8-Br-cADPR suppresses degeneration by antagonizing cADPR signaling downstream of Sarm1. We also tested 8-Br-7-CH-cADPR, which is a more potent cADPR antagonist than 8-Br-cADPR (Moreau et al., 2011). Axonal treatment with 8-Br-7- CH-cADPR significantly decreased paclitaxel-induced degeneration at the concentration as low as 0.1µM (FigureS3C), further indicating cADPR is involved in paclitaxel-induced axon degeneration. One alternative explanation is that 8-Br-cADPR may function as a direct inhibitor of Sarm1 and suppresses cADPR production. We performed an *in vitro* Sarm1 activity assay using purified Sarm1 protein supplemented with NAD/NMN and found that 8-Br-cADPR did not affect Sarm1 NADase activity as assayed by the production of ADPR or nicotinamide (Nam) (FigureS3D-E). Moreover, axonal treatment with 8-Br-cADPR did not alter sTIR dimerization-induced cADPR production (FigureS2B), indicating that the neuroprotective effect of 8-Br-cADPR is downstream of Sarm1. These results collectively demonstrate that cADPR-dependent calcium signaling contributes to paclitaxel-induced axon degeneration *in vitro*.

**Figure3.**
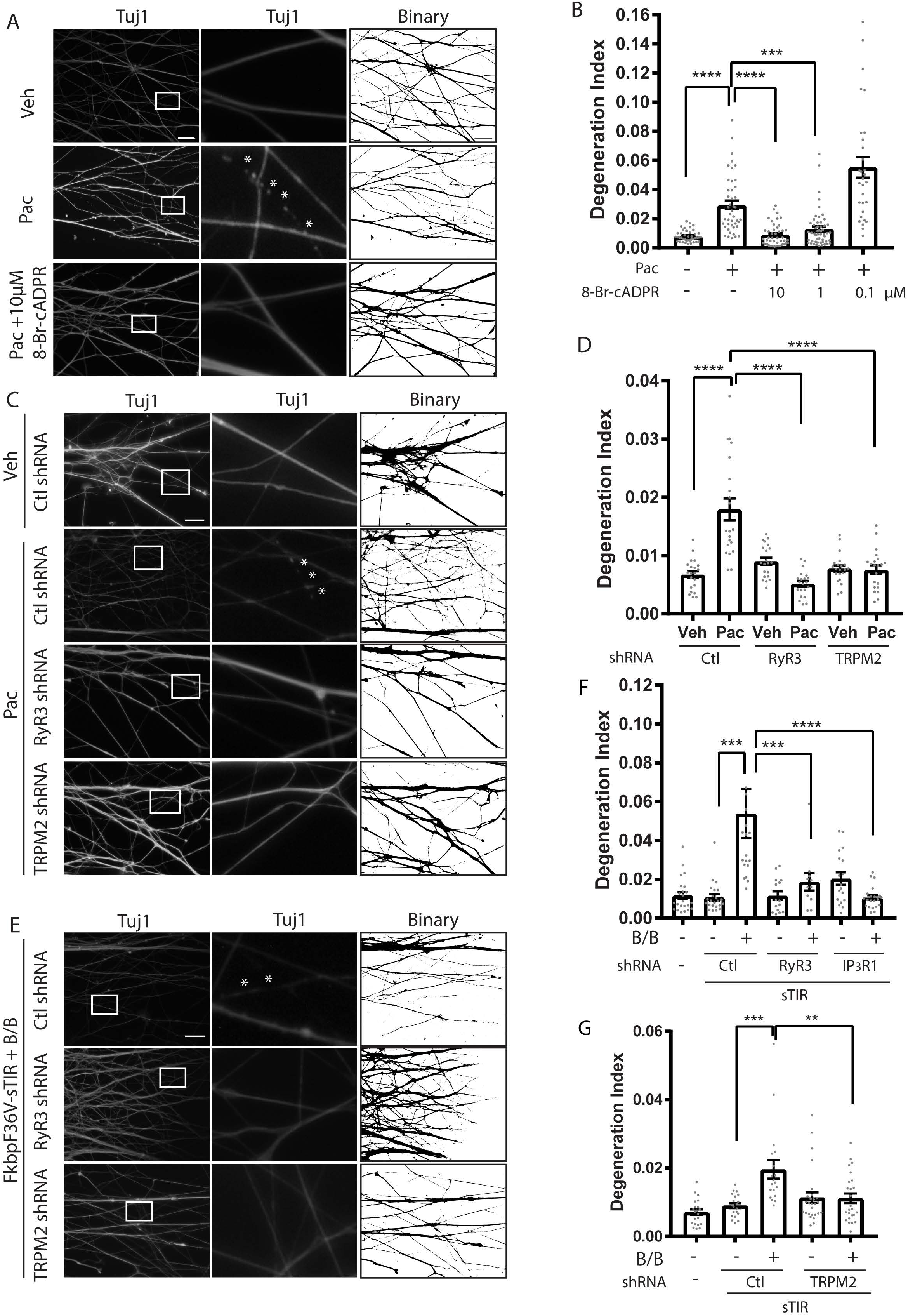
**Paclitaxel-induced axon degeneration is cADPR dependent.** (A) Tuj1 immunostaining and corresponding binarized images of axons in compartmentalized cultures after 24hrs of 30nM paclitaxel (Pac), 30nM paclitaxel with 10µM 8-Br-cADPR or DMSO (Veh) added to axons. White boxes outline regions shown at higher magnification in the center panels. White stars indicate fragmented region of axons displayed as interruptions in Tuj1 continuity. Scale bar: 20µm. (B) Degeneration index of (A). Data represent mean +/- SEM; individual data points are shown. N represents number of images. Data pooled from seven independent experiments. ****p < 0.0001 by one-way ANOVA with Tukey’s multiple comparison test. (C) Tuj1 immunostaining and corresponding binarized images of axons after 24hrs of 30nM paclitaxel (Pac) or DMSO (Veh) added to axons following lentiviral infection with RyR3 or TRPM2, or control shRNA (Ctl). White boxes outline regions shown at higher magnification in the center panels. White stars indicate fragmented region of axons displayed as interruptions in Tuj1 continuity. Scale bar: 20µm. (D) Degeneration index of (C). Data represent mean +/- SEM. Individual data points are shown. N represents number of images. Data pooled from three independent experiments. **p<0.01 ***p<0.001 ****p < 0.0001 by one-way ANOVA with Tukey’s multiple comparison test. (E) Tuj1 immunostaining and corresponding binarized images of axons of DRG neurons co-expressing FkbpF36V-sTIR and indicated shRNA, 3hrs after 50nM B/B homodimerizer added to axons. White boxes outline regions shown at higher magnification in the center panels. White stars indicate fragmented region of axons displayed as interruptions in Tuj1 continuity. Scale bar: 20µm. (F-G) Degeneration index of (E). Data represent mean +/- SEM. Individual data points are shown. N represents number of images. Data pooled from three independent experiments. Two data points 0.199, 0.208 of Ctl + sTIR + B/B fall outside y-axis limit not shown in (F). **p<0.01 ***p<0.001 ****p < 0.0001 by one-way ANOVA with Tukey’s multiple comparison test.

cADPR modulates intracellular calcium release through intracellular RyRs on the endoplasmic reticulum (Galione et al., 1991; Guse, 1999; Meszaros et al., 1993; Sonnleitner et al., 1998). As RyR3 is the major type of RyRs in DRG neurons (Lokuta et al., 2002), we knocked down RyR3 using lentivirus-delivered shRNA (FigureS3F) and found that RyR3 depletion inhibited paclitaxel-induced axon degeneration (Figure3C-D). These data indicate that RyR3-dependent calcium release from the ER contributes to paclitaxel-induced degeneration. cADPR also directly binds to and activates TRPM2, a calcium-permeable, non-selective cation channel predominantly localized on the plasma membrane (Kolisek et al., 2005; Yu et al., 2019). Knockdown of TRPM2 also protects against paclitaxel-induced axon degeneration in DRG cultures (Figure3C-D, FigureS3G). These results suggest that calcium release from both intra- and extracellular sources are required for axon degeneration downstream of paclitaxel.

To test whether RyR3 and TRPM2 open in response to activation of Sarm1, we co-expressed FkbpF36V- sTIR and shRNA targeting RyR3 or TRPM2 individually in DRG neurons. Knockdown of RyR3 or TRPM2 individually inhibited axon degeneration caused by sTIR dimerization (Figure3E,F,G). As a control, dimerization of FkbpF36V-mTIR did not lead to axon degeneration (FigureS3H). While cADPR-dependent calcium mobilization is known to be IP_3_R independent, knockdown of IP_3_R1 also suppressed axon degeneration caused by FkbpF36V-sTIR dimerization (Figure3F, FigureS3I). Together these results suggest that calcium signaling pathways independent of cADPR also contribute to Sarm1 activation-induced degeneration, consistent with the idea that cADPR synergizes with other factors to modulate calcium release and cause degeneration. These data also indicate that both ER calcium stores and extracellular calcium are required for paclitaxel-induced intra-axonal calcium influx and the ensuing axon degeneration.

Sarm1 activation also produces ADPR, the hydrolyzed form cADPR. Up to 20% of commercially available 8-Br-cADPR may consist of 8-Br-ADPR, an antagonist of ADPR (personal communication, Timothy Walseth). As ADPR is a more potent ligand for TRPM2 than cADPR (Huang et al., 2018; Kolisek et al., 2005; Yu et al., 2019), we asked whether both components may contribute to the observed efficacy. We treated DRG axons with 8-Br-ADPR and found 8-Br-ADPR also suppressed paclitaxel-induced axon degeneration (Figure S3J), consistent with our finding that TRPM2 and RyR both contribute to paclitaxel-induced degeneration.

Calcium dysregulation is also implicated in Sarm1-dependent axon degeneration in response to physical injury. We then asked whether an antagonist of cADPR also protect against Wallerian degeneration. Interestingly, we found 8-Br-cADPR does not prevent axotomy or mitochondrial dysfunction induced axon degeneration (FigureS3K-L). Our data indicate that there are qualitative and/or quantitative differences in the function of cADPR between axon degeneration caused by axotomy and the chemotherapeutic agent paclitaxel.

### Antagonizing cADPR shows neuroprotective effects against paclitaxel *in vivo*

Our results indicate that Sarm1 and cADPR are critical for paclitaxel-induced axon degeneration *in vitro*. To assess the roles of cADPR in paclitaxel-induced peripheral neuropathy *in vivo*, we treated 2-month-old C57BL6/J mice with multiple intraperitoneal injections of paclitaxel. Animals were also treated with 8-Br-cADPR or saline control. We then evaluated the paclitaxel-induced neuropathy *in vivo* at both behavioral and pathological levels (Figure4A). A key feature of CIPN in patients is allodynia, a condition in which pain is caused by stimuli that do not normally elicit pain. To assess allodynia in mice before and after paclitaxel, we compared the sensitivity of the mice to mechanical stimuli using filaments of increasing rigidity (Von Frey Filament tests). A much smaller force triggered a withdrawal response in animals after paclitaxel treatment, while vehicle-treated control mice showed no difference before and after treatment (Figure4B). We found that systemic treatment with 8-Br-cADPR significantly suppressed paclitaxel-induced excess pain sensitivity (Figure4B). Loss of intra-epidermal nerve fiber (IENF) density is a typical pathological feature of CIPN consistently observed in both patients and animal models (Han and Smith, 2013). We collected footpads from the mice after the final behavior test and examined IENF of both thin (non-dermal papillae-containing) and thick (dermal papillae-containing) skin. Paclitaxel treatment significantly decreased IENF density in both thin (Figure4C-D) and thick (Figure4C,E) skin compared to vehicle-treated mice. Treatment with 8-Br-cADPR provided partial protection from paclitaxel-induced loss of innervation (Figure4C-E). There were no significant body weight changes before and after treatment with either paclitaxel alone or paclitaxel together with 8-Br-cADPR (FigureS4), indicating minimal non-specific toxicity of these treatments. These results demonstrate a neuroprotective effect of 8-Br-cADPR against paclitaxel-induced peripheral neuropathy *in vivo*, suggesting that therapies targeting cADPR may be an effective strategy for CIPN.

**Figure4.**
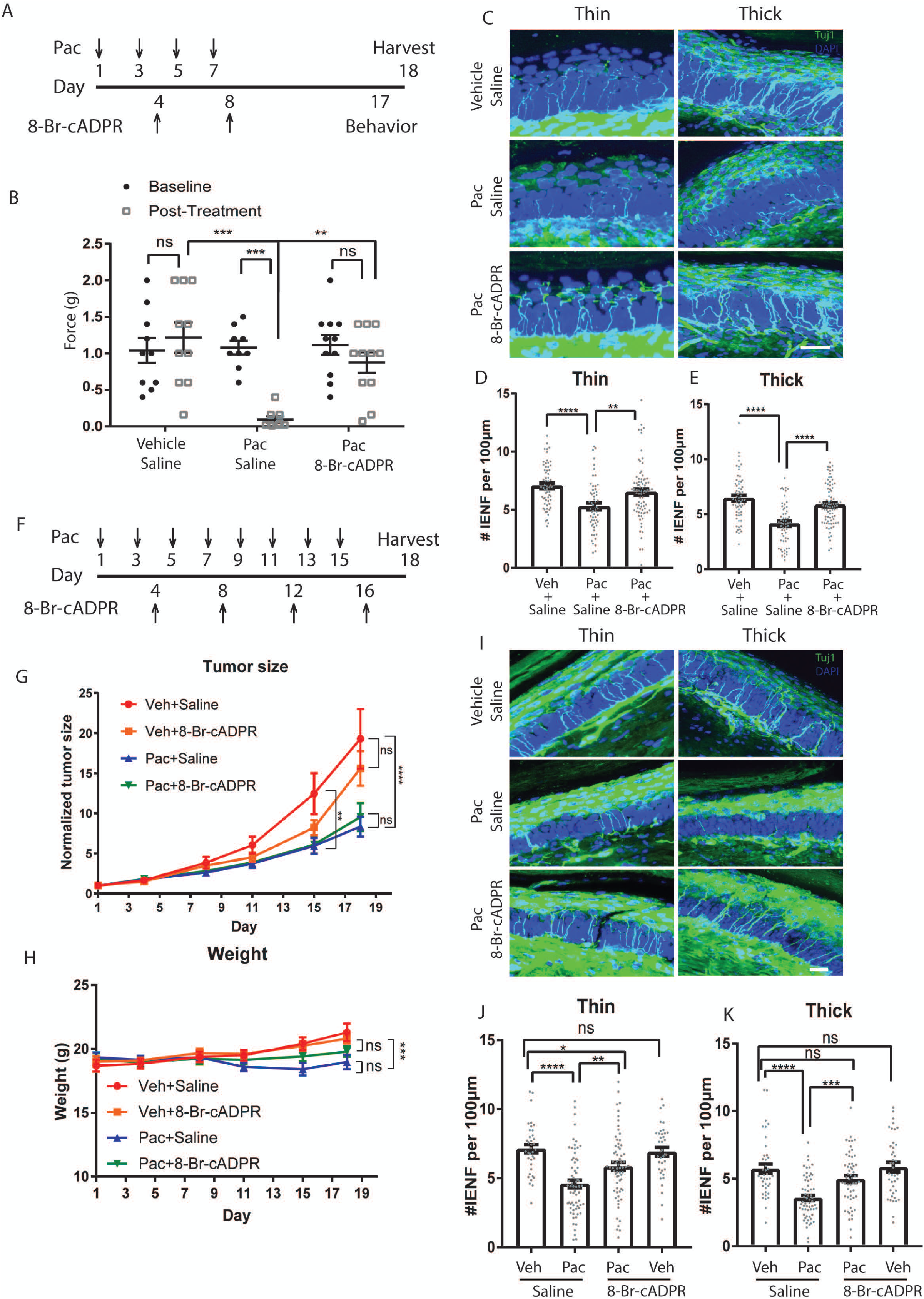
**cADPR antagonist protects against paclitaxel-induced peripheral neuropathy *in vivo*** (A) Schematic of experimental design. Paclitaxel (20mg/kg) or vehicle was injected intraperitoneally on days 1, 3, 5 and 7, and 8-Br-cADPR (2mg/kg) or saline was injected intraperitoneally on days 4 and 8. Final behavior test with Von Frey Filaments was performed on day 17 and mice were euthanized for IENF analysis on day 18. (B) Mechanical pain threshold assessments of mice using Von Frey filaments before (Baseline) and 10 days after the final injection (Post-Treatment) of paclitaxel or vehicle control. Data represent mean +/- SEM; individual data points are shown. N represents number of animals. **p < 0.01 ***p<0.001 by two-way ANOVA with Tukey’s multiple comparison test. (C–E) Quantification and representative images of Tuj1-positive sensory fibers (green) entering the epidermis per 100µm epidermal length in thin (C left and D) and thick (C right and E) skin. DAPI counterstain (blue); scale bar: 20µm. **p < 0.01 ***p<0.001 ****p<0.0001 by one-way ANOVA with Tukey’s multiple comparison test. Data represent mean +/- SEM; individual data points are shown. N represents number of images; 6-10 images from each animal. (F) Schematic illustration of experimental design. Paclitaxel or vehicle was injected intraperitoneally on day 1, 3, 5, 7, 9 11, 13 and 15, and 8-Br-cADPR or saline was injected intraperitoneally on day 4, 8, 12 and 16. Mice were euthanized for IENF analysis on day 18. (G) Normalized tumor size of mice with E0771 breast cancer cells treated with vehicle + saline, paclitaxel + saline, paclitaxel + 8-Br-cADPR or vehicle + 8-Br-cADPR. Data represents mean +/- SEM; N represents number of tumor. 5 animals with bilateral tumors were used for each condition. **p < 0.01 ****p<0.0001 by two-way ANOVA with Tukey’s multiple comparison test at each time point. (H) Body weights of tumor-bearing mice treated with vehicle + saline, paclitaxel + saline, paclitaxel + 8- Br-cADPR or vehicle + 8-Br-cADPR. Data represent mean +/- SEM; N represents number of animals. 5 animals were used for each condition. ***p < 0.01 by two-way ANOVA with Tukey’s multiple comparison test at each time point. (I-K) Representative images and quantification of Tuj1-positive sensory fibers (green) entering the epidermis per 100µm epidermal length in thin (I left and J) and thick (I right and K) skin. DAPI counterstain (blue). Scale bar: 20µm. Data represent mean +/- SEM; individual data points are shown. N represents number of images. 8-16 images from each animal. * p<0.05 **p < 0.01 ***p<0.001 ****p<0.0001 by one-way ANOVA with Tukey’s multiple comparison test.

A concern in targeting cADPR for therapeutic purposes is the possibility that this intervention might decrease the anti-neoplastic efficacy of paclitaxel, as calcium homeostasis is also involved in cell cycle regulation and tumor progression (Stewart et al., 2015; Yang et al., 2020). To address this, we used an immunocompetent breast cancer mouse model using the breast cancer line E0771 in C57BL6/J mice. We implanted E0771 tumor cells orthotopically into the mammary fat pads of 7-week-old C57BL6/J mice. We began paclitaxel injections one week after tumor cell injection, when the tumor size reached approximately 5mm in diameter. To maximize the effect of paclitaxel on tumor growth, paclitaxel or vehicle was injected every other day for a total of 8 injections. 8-Br-cADPR or saline was injected after every two paclitaxel or vehicle injections (Figure4F). Tumor sizes increased gradually in mice treated with vehicle + saline up to day 18 (Figure4G), at which point some of these mice approached humane endpoint. Treatment with 8-Br-cADPR alone did not alter tumor growth (Figure4G). Paclitaxel treatment significantly inhibited tumor growth (Figure4G), and combined treatment with paclitaxel + 8-Br-cADPR did not affect the ability of paclitaxel to suppress tumor growth (Figure4G), indicating that 8-Br-cADPR does not interfere with the therapeutic efficacy of paclitaxel. Extended treatment with paclitaxel led to a small but statistically significant weight decrease, while 8-Br-cADPR did not cause weight changes when injected alone or with paclitaxel (Figure4H). We then further analyzed CIPN in these mice by evaluating IENF innervation. IENF density of both thin and thick skins was significantly decreased in paclitaxel-treated mice compared to vehicle control, and 8-Br-cADPR partially but significantly rescued the IENF density loss in both thin and thick skins (Figure4I-K). Notably, 8-Br-cADPR treatment alone did not affect IENF density (Figure4J-K). Together, our data demonstrate that 8-Br-cADPR protects against paclitaxel-induced allodynia and nerve fiber degeneration *in vivo* without affecting the anti-tumor effect of paclitaxel, indicating it may provide a good therapeutic approach to CIPN.

## DISCUSSION

### cADPR as a calcium modulator in paclitaxel-induced axon degeneration

Understanding the role of calcium dysregulation in chemotherapy-induced axon degeneration requires direct monitoring of axonal calcium flux following drug treatment. In this study, we directly visualized and analyzed axonal calcium flux after paclitaxel treatment and demonstrated that local treatment of paclitaxel causes a progressive increase of axonal calcium. While previous studies have focused on cADPR as a readout of Sarm1 activity (Sasaki et al., 2020), here we show that paclitaxel-induced axonal calcium flux and degeneration is partially attributed to cADPR produced by Sarm1 activity. cADPR functions as a calcium modulator of both RyR and TRPM2, and we find that multiple calcium channels, including RyR3, TRPM2 and IP3R1, all contribute to axon degeneration initiated by paclitaxel or Sarm1 activation. Thus, multiple intra- and extracellular sources of calcium are involved in axonal calcium flux caused by paclitaxel. Our studies suggest a model wherein cADPR produced by activated Sarm1 contributes to the axonal calcium flux by modulating calcium-induced calcium release.

In contrast to the efficacy of cADPR antagonists in preventing CIPN both *in vitro* and *in vivo*, we found that a cADPR antagonist was unable to prevent axotomy-induced axon degeneration. Our data indicate that although axonal degeneration triggered by paclitaxel and axotomy are both Sarm1-dependent, the two signaling cascades are functionally different. We previously showed paclitaxel also causes axonal depletion of Bclw, which leads to IP_3_R1 dependent calcium dysregulation. Thus multiple factors contribute to changes in calcium regulation and the degeneration process in CIPN. Further analyses will be required to identify the mechanisms that differentiate slow-progressing Sarm1 dependent degeneration caused by paclitaxel from the more acute and severe perturbation of axotomy.

### Sarm1 activation downstream of paclitaxel

The mechanisms that activate Sarm1 are just beginning to be defined. A recent study demonstrates that Sarm1 is a metabolic sensor that can be allosterically regulated by the ratio of NMN and NAD (Figley et al., 2021). Although we do not yet know how paclitaxel activates Sarm1, an attractive idea is that paclitaxel causes defective axon transport of NMNAT2, a highly labile protein that converts NMN to NAD, and must be continuously provided to the axon by fast axonal transport (Gilley and Coleman, 2010). Indeed depletion of the axon survival factor NMNAT2 leads to Sarm1 dependent axon degeneration (Gilley et al., 2015). We speculate that microtubule-dependent transport of NMNAT2 may be disrupted by paclitaxel, which leads to NMNAT2 depletion, changes in the ratio of NMN and NAD and Sarm1 activation. Other molecules including mitochondrial antiviral signaling protein (MAVS) and syndecan-2 (Sdc2) have been suggested as additional potential regulators of Sarm1 activity (Chen et al., 2011; Mukherjee et al., 2013). Thus, it is also possible that transport of MAVS and/or Sdc2 may be impaired with paclitaxel treatment, leading to changes in axonal MAVS and/or Sdc2 that further activate Sarm1.

### Tumor mouse model in CIPN research and therapeutic implication of cADPR

Our *in vivo* studies corroborate the involvement of cADPR in paclitaxel-induced degeneration. Paclitaxel-treated mice exhibit heightened pain sensitivity and loss of intra-epidermal nerve endings, recapitulating the allodynia and dying back nerve degeneration observed in patients. Intraperitoneal administration of 8-Br-cADPR attenuates paclitaxel-induced peripheral neuropathy both behaviorally and pathologically. The robust protection of 8-Br-cADPR *in vivo*, however, may involve other components besides the nerve itself since the role of cADPR as a calcium modulator has also been implicated in the immune system and glial cells (Banerjee et al., 2008; Guse et al., 1999; Partida-Sanchez et al., 2001). Nevertheless, our results raise the possibility of using cADPR as a potential therapeutic target for treating CIPN.

Preventative treatments of CIPN will be given to patients who are undergoing treatment for a malignant cancer. Therefore, in searching for new therapies for CIPN, it is critical that the neuropathic and anti-neoplastic effects of the microtubule-targeting chemo-drugs can be uncoupled from one another. Using an immune competent mouse model of breast cancer, we were able to examine both peripheral neuropathy progression and tumor growth in the same animals. Consistent with prior studies, we found that tumor growth was significantly inhibited by paclitaxel in this model (Bourgeois-Daigneault et al., 2016); and these paclitaxel-treated mice also exhibited decreased IENF density, confirming that paclitaxel causes peripheral neuropathy in this model. Treatment with the cADPR competitive antagonist, 8-Br-cADPR, improved the paclitaxel-induced neuropathy without interfering with its anti-tumor effect. Thus, the effects of paclitaxel on cancer growth can be disentangled from its effects on nerve innervation. Our results *in vitro* and *in vivo* collectively indicate the potential of targeting cADPR as a novel and promising therapeutic approach for preventing CIPN. Further studies investigating the ability of new small molecule inhibitors of Sarm1 (Hughes et al., 2021) to prevent or delay CIPN *in vivo* without altering therapeutic effects of chemotherapies may provide additional therapeutic approaches to this major cause of poor quality of life in cancer survivors.

## MATERIALS AND METHODS

### Mouse line and animal care

All experimental procedures were conducted in accordance with the National Institutes of Health guidelines and were approved by the Dana-Farber Cancer Institute or the Washington University School of Medicine, St. Louis Institutional Animal Care and Use Committee.

Timed pregnant Sprague-Dawley rats and CD1 mice were purchased from Charles River. C57BL6/J mice were purchased from Jackson Laboratory.

### DRG neuron culture

DRG were dissected from E15 rat embryos, dissociated and plated in Matrigel (1:45; Thermo Fisher)- coated p35 dishes, Campenot device or poly-D-lysine/Laminin-coated microfluidic chambers. DRG cultures were maintained in NeuroBasal medium supplemented with 2% B27, 1% Glutamax, 1% penicillin and streptomycin, 0.08% glucose, 1-100ng/mL NGF/BDNF (PeproTech) and 0.5µM Cytarabine (AraC). Cultures were maintained in incubators at 37°C with 7.5% CO_2_. For non-compartmentalized mass culture, 300,000 cells were plated in each p35 dish. BDNF + NGF were added at a concentration of 100ng/mL for 2 days and reduced to 10ng/mL for 5-6 days. Campenot device were prepared as described (Fenstermacher et al., 2015). BDNF + NGF were added to the cell body compartment at a concentration of 10ng/mL and to the axon compartment at a concentration of 100ng/mL for 2 days. On day 5 neurotrophins were removed from the cell body compartment and reduced to 1ng/mL in the axon compartments for 2-3 days. Paclitaxel (30nM) or DMSO (0.0025%) was added to the axon chambers on DIV7-8 for 24 hours before fixation. For TIR domain dimerization, 50nM B/B homodimerizer was added to the axon chambers for 3 hours before fixation. For degeneration assay with 8-Br-cADPR treatment, 8- Br-cADPR (0.1-10µM) was added to axon chambers 1hr before paclitaxel or B/B homodimerizer addition and continued to be present in the axon chambers with paclitaxel or B/B homodimerizer until fixation.

Microfluidic chambers (Xona Microfluidics) were prepared according to manufacturer’s instruction with laminin and poly-D-Lysine coating. 30,000 neurons were plated in the cell body compartment. BDNF + NGF were added to the cell body compartment at 50ng/mL and to the axon compartment at 100ng/mL for 1 day, and reduced to 10ng/ml (cell bodies) and 100ng/ml (axons) for 1 day, and maintained in 5ng/ml (cell bodies) and 10ng/ml (axons) for 3-5 days. Paclitaxel (600nM) or 0.5% DMSO was added to axon compartments on DIV4-5 for 48hrs. For TIR domain dimerization, 50nM B/B homodimerizer was added to the axon chambers on DIV4-5 for 4 hours. For experiment with 8-Br-cADPR, 8-Br-cADPR was added to axon chamber 1hr before paclitaxel or 50nM B/B homodimerizer and continued to be present in axon together with paclitaxel or B/B homodimerizer.

For axotomy induced axon degeneration, mouse dorsal root ganglion (DRG) culture was performed as described previously (Sasaki et al., 2016). Briefly, DRG was dissected from embryonic days 13.5-14.5 CD1 mouse (Charles River Laboratories) and incubated with 0.05% Trypsin solution at 37 °C for 20 minutes. Then, cell suspensions were triturated by gentle pipetting and washed with Neurobasal culture medium (Gibco) containing 2% B27 (Invitrogen), 50 ng/ml nerve growth factor (Envigo Bioproducts, Cat. #B5017), 1µM uridine (Sigma), 1µM 5-fluoro-2’-deoxyuridine (Sigma), penicillin and streptomycin. Cell suspensions were placed in the center of the well using 24-well tissue culture plate (Corning) coated with poly-D-Lysine (0.1 mg/ml; Sigma) and laminin (3 µg/ml; Invitrogen).

### LC-MS/MS analysis

Metabolites extraction was done following nucleotide extraction protocol provided by McGill metabolic core facility. Briefly, both mass culture and Campenot culture were washed with ice-cold 150mM ammonium formate buffer and harvested in ice-cold 50%/50% (v/v) methanol/LC-MS grade water, followed by three solvent extraction (50% methanol: acetonitrile: dichloromethane: water = 380µl: 220µl: 600µl: 300µl). The aqueous phase was then isolated and dried using vacuum centrifugation. For semi-quantitative targeted metabolite analysis of nucleotides, samples were injected onto an Agilent 6470 Triple Quadrupole (Agilent Technologies, Santa Clara, CA, USA). Chromatography was achieved using a 1290 Infinity ultra-performance LC system (Agilent Technologies, Santa Clara, CA, USA) Separation was performed on a Scherzo SM-C18 column 3μm, 3.0×150mm (Imtakt Corp, JAPAN) maintained at 10°C. The chromatographic gradient started at 100% mobile phase A (5mM ammonium acetate in water) with a 5 min gradient to 100% B (200 mM ammonium acetate in 20% ACN / 80% water) at a flow rate of 0.4ml/min. This was followed by a 5 min hold time at 100% mobile phase B and a subsequent re-equilibration time (6 min) before next injection. Samples were individually re-suspended in 30μL of cold water and 5μL volume was injected immediately after sample preparation to help preserve unstable metabolites.

Multiple reaction monitoring (MRM) transitions were optimized on standards for each metabolite measured. MRM transitions and retention time windows are summarized in Table1. An Agilent JetStreamTM electro-spray ionization source was used in positive ionization mode with a gas temperature and flow were set at 300°C and 5 L/min respectively, nebulizer pressure was set at 45 psi and capillary voltage was set at 3500V. Relative concentrations were determined from external calibration curves prepared in water. Ion suppression artifacts were not corrected; thus, the presented metabolite levels are relative to the external calibration curves and should not be considered as absolute concentrations. Data were analyzed using MassHunter Quant (Agilent Technologies). The amount of each metabolite per sample was normalized to cell number.

**Table1:** Summary of MRM transitions and retention time windows

| Compound Name | Precursor Ion | Product Ion | Ret Time (min) | Fragmentor | Collision Energy | Polarity |
| --- | --- | --- | --- | --- | --- | --- |
| ADPR | 560.1 | 97.14 | 4.47 | 166 | 64 | Positive |
| ADPR | 560.1 | 136 | 4.47 | 166 | 36 | Positive |
| ADPR | 560.1 | 347.9 | 4.47 | 166 | 25 | Positive |
| ADPR | 560.1 | 427.8 | 4.47 | 166 | 25 | Positive |
| cADPR | 542.1 | 97.1 | 3.60 | 166 | 64 | Positive |
| cADPR | 542.1 | 136 | 3.60 | 166 | 36 | Positive |
| NAD | 664.1 | 136.1 | 4.55 | 100 | 56 | Positive |
| NAD | 664.1 | 428 | 4.55 | 100 | 24 | Positive |
| NADP | 744.1 | 136.3 | 4.69 | 120 | 56 | Positive |
| NADP | 744.1 | 604 | 4.69 | 120 | 12 | Positive |

### Plasmids construction

Lentiviral constructs expressing shRNAs targeting rat:

Sarm1 (CAGGTAGATGGTGATTTGCTT)

RyR3 (AAGCTCCTGACAAATCACTAT)

CD38 (GAGCATCCATCATGTAGACTT)

were generated using pLKO.1 vector following instructions provided by Addgene. The shRNA sequence was selected using BLOCK-iT™ RNAi Designer provided by Thermo Fisher Scientific. Lentiviral constructs expressing shRNA targeting rat IP_3_R1 (TRCN0000321161; Sigma-Aldrich) and TRPM2 (TRCN0000068488 Sigma-Aldrich) were purchased from Sigma-Aldrich. Lentiviral particles were then generated and knockdown efficacy was verified by reduction of protein or mRNA levels in mass cultured DRGs (FigureS1A-B, FigureS3F-G, I). TurboGFP-targeting shRNA (SHC004 Sigma-Aldrich) and a universal non-targeting shRNA (LV015-G ABM) were used as control shRNAs.

Lentiviral constructs expressing FkbpF36V-Sarm1-TIR and FkbpF36V-Myd88-TIR were generated as described (Gerdts et al., 2015). For FCIV-hSarm1, a full-length isoform of the canonical 724 amino acid human SARM1 was introduced into lentivirus vector FCIV using the Takara HD InFusion Cloning Kit and complimentary oligonucleotides. The expressed SARM1 protein contains optimized start codons K3, I4, H5 and is fused to 2 X Strep Tag at the N-terminus. The absence of mutations and other PCR errors was confirmed by sequencing (Genewiz).

### Lentivirus generation and transduction

Lentivirus was generated following protocols provided by Addgene. Briefly, transfer plasmid, packaging plasmid pCMV-dR8.91 and envelop plasmid pCMV-VSV-G were transfected into HEK293T cells using FUGENE6 or Lipofectamine2000 following manufacturer’s instruction. The media containing virus were harvested 48 and 72hr after transfection and pooled, and then concentrated to approximately 1/12 of the original volume using Amicon Ultra-centrifugal filter. For lentivirus infection of DRG neurons in compartmented Campenot cultures, 25-50µl lentivirus was added to the middle chamber of Campenot culture containing cell bodies on DIV2. For microfluidic culture, 100µl lentivirus of TIR domains or 25µl lentivirus of shRNA was added to the cell body chamber on DIV1.

### Quantitative reverse transcription-PCR

RNA was extracted from DRG neurons using TRIzol (Invitrogen) according to manufacturer’s instruction. Reverse transcription was performed using the SuperscriptIII first strand synthesis (Thermo Fisher) according to the manufacturer’s instruction and quantitative real-time PCR was performed using Taqman Gene expression assays (Applied Biosystems) to analyze expression of SARM1 (Rn01750947_m1), TRPM2 (Rn01429410_m1), CD38 (Rn00565538_m1), RyR3(Rn01486097_m1). Data were normalized to GAPDH (Applied Biosystems) for each sample.

### Western Blot

For assessing IP_3_R1 level, DRG neurons transduced with IP_3_R1 or control shRNA for 6-7 days were lysed with non-ionic detergent. Lysates were separated by 3%–8% Tris-Acetate SDS-Page (Thermo fisher) and probed against rabbit IP_3_R1 (1:1000; Thermo Fisher) and rabbit pan-actin (1:1000; Cell signaling). Bands were visualized with secondary antibodies conjugated to HRP (1:10,000; Bio-Rad) and SuperSignal chemiluminescent substrates signal (Thermo Fisher). Blots were imaged using AI600 Chemiluminescent Imager (GE Healthcare).

### Axonal degeneration assay

Neurons in Campenot cultures were fixed at room temperature with 4% PFA diluted 1:2 in media for 10 min, then undiluted 4% PFA for an additional 20 min. Cultures were permeabilized and blocked in 3% BSA and 0.1% Triton X-100 for 1 hour at room temperature, and incubated with mouse anti-Tuj1 (1:500; BioLegend) overnight at 4°C. Cultures were then washed twice with PBS and incubated with goat anti-mouse AlexaFluor (1:400; Invitrogen) and DAPI for 1 hour at room temperature. Campenot dividers were then removed and samples were mounted with Fluoromount-G (Southern Biotech 0100-01). Images of distal axon tips were obtained using a 40X air objective; the images were binarized, and axonal degeneration was quantified as degeneration index ratio of fragmented axons divided by total axon area in each image (Pease-Raissi et al., 2017; Sasaki et al., 2009).

Neurons in microfluidic cultures, were permeabilized and blocked in 3% BSA and 0.1% Triton X-100 for 1 hour at room temperature, and incubated with mouse anti-Tuj1 (1:500; BioLegend) overnight at 4°C. Cultures were then washed twice with PBS and incubated with donkey anti-mouse AlexaFluor (1:400; Invitrogen) and DAPI for 1 hour at room temperature. Teflon dividers were then removed and samples were mounted with Fluoromount-G (Southern Biotech 0100-01). Ten to twenty fields of axons for each sample were imaged with a Nikon Ti-E at 37°C (60x oil 1.4NA objective) with 7.5% CO_2_. The same XY positions were imaged for treatments and control for each experiment. Images were binarized and axon area was measured using NIH ImageJ software. Data was then normalized to DMSO-treated control.

### Axotomy induced axon degeneration assay

Mouse DRG neurons were placed in the center of the well using 24-well tissue culture plate. On DIV7, 1 or 10µM 8-Br-cADPR was pre-incubated 30 mins before axon injury. Axons from DRG neurons were either transected using a microsurgical blade under a microscope or applied with 50µM CCCP (carbonyl cyanide m-chlorophenylhydrazone) for axon degeneration assay. Bright-field images of distal axons (20 fields/ well and 3wells/ conditions) were acquired at 0-24 hours after axon injury using a high content imager (Operetta; PerkinElmer) with a 20x objective lens. Axon degeneration was quantified using degeneration index calculated with ImageJ.

### Calcium imaging

E15 DRG neurons in microfluidic devices were transduced on DIV1 with 5-10µl AAV9-axonal-GCaMP6s- P2A-mRuby3 (Broussard et al., 2018) (AAV9 was generated by Applied Biological Materials, Canada) alone or together with 25µl lentivirus expressing shRNA targeting tGFP, Sarm1 or CD38 for 24 hours. Cultures were maintained in phenol red-free NeuroBasal medium supplemented with 2% B27, 1% Glutamax, 1% penicillin and streptomycin, 0.08% glucose, NGF/BDNF and 0.5µM Cytarabine (AraC) for 4-5 days before imaging. For neurons transduced with lentivirus expressing shRNA, 1µg/mL puromycin was added to the cell body on DIV3 for 2 days for selection and was removed before imaging. Eight to ten fields of axon chamber for each sample were imaged with a Nikon Ti-E at 37°C (60X oil 1.4NA objective) with 7.5% CO_2_ before treatment as time zero. Then, 600nM paclitaxel or 0.5% DMSO was added to axon compartments. Cultures were maintained in incubators at 37°C with 7.5% CO_2_. The same fields of axons for each sample were imaged manually at 12, 16, 20, 24, 36, 40, 44 and 48 hours after treatment. For experiments with 8-Br-cADPR, 20µM 8-Br-cADPR was added to axon chamber 1 hour before paclitaxel and continued to be present in axon together with paclitaxel.

For FkbpF36V-sTIR-induced calcium flux, DIV1 DRG neurons were transduced with 5-10µl AAV9-axonal-GCaMP6s-P2A-mRuby3 together with 100µl lentivirus expressing FkbpF36V-sTIR or FkbpF36V-mTIR. Cultures were maintained in phenol red-free NeuroBasal medium supplemented with 2% B27, 1% Glutamax, 1% Pen/Strep, 0.08% glucose, NGF/BDNF and 0.5 µM AraC for 4-5 days before imaging. Expressions of FkbpF36V-sTIR or mTIR were verified based on Cerulean expression. Eight fields of axon chamber for each sample were imaged before treatment and every 1 hour after addition of 50nM B/B homodimerizer to the axon chamber. For experiments with 8-Br-cADPR, 10µM 8-Br-cADPR was added to axon chamber 1 hour before 50nM B/B homodimerizer was added and continued to be present in axon chamber.

Both GCaMP6s and mRuby3 signals were acquired at each time point using the Nikon Ti-E at 37°C (60X oil 1.4NA objective) with 7.5% CO_2_. Images were analyzed using NIH ImageJ software. For each field at one time point, a mask was generated from mRuby3 channel, background fluorescence was subtracted, and GCaMP6s and mRuby3 fluorescence intensities were measured. The GCaMP6s for each field at each time point was normalized to the mRuby3 intensity. The change in fluorescence was measured per field of axons according to the equation [(GCaMP6s/mRuby)/(GCaMP6s/mRuby)_0_], where (GCaMP6s/mRuby)_0_ is the baseline at time 0 before treatment. Treatments and the corresponding controls were done in parallel for each independent experiment.

### Expression and purification of Sarm1 protein

Human SARM1 was transfected to 150mm diameter cell culture dish with 50% confluence of HEK293T cells. 15µg of plasmid was mixed with 75µg of polyethylenimine (PEI, 1mg/mL, pH 7.0) and transfected to the cells. Cells were harvested 48 hours after transfection and re-suspended in 100mM Tris-Cl, pH 8.0, and 150mM NaCl with Protease Inhibitor Cocktail (Pierce) before lysed with sonication on ice. After centrifugation at 18,000xg for 10 minutes and 3 times to remove the cell debris, supernatant was mixed with PureCube HiCap Streptactin MagBeads (Cube Biotech) for 1 hour. After washing 3 times with 25 mM HEPES, pH 7.5, and 150mM NaCl, SARM1-laden beads were stored in the same buffer plus 1mM TCEP and stored at -80°C. Protein purify was assessed by 4-12% SDS-PAGE by Coomassie staining. Protein concentration was determined using ImageJ (NIH) against BSA standard with known concentrations running on the same SDS-PAGE.

### Sarm1 *in vitro* activity assay

Sarm1 *in vitro* activity was assessed as described previously with modification (Essuman et al., 2017) . Briefly, human SARM1 (15nM)-laden beads were mixed with various concentrations of 8-Br-cADPP in 50mM HEPES, pH 7.5, 50µM NAD, and 25µM NMN at 25 °C. Reaction was carried out in a ThermoMixer. At various time points, the reaction was stopped by taken 50µL from the reaction mixture and mixing with 50µL 0.5M perchloric acid (HClO4) before placing on ice for 10 minutes. After centrifugation at 18,000xg for 10 minutes, supernatant was mixed with 6µL 3M K2CO3 for neutralization. Samples were placed on ice for another 10 minutes and centrifuged one more time. 45µL of supernatant containing extracted metabolites was mixed with 5µL 0.5M Potassium Phosphate buffer and quantified by HPLC (Nexera X2) with Kinetex (100 × 3 mm, 2.6µm; Phenomenex) column. ADPR and Nam production rates were calculated from samples taken at various time points at each 8-Br-cADPR concentration.

### Paclitaxel treatment *in vivo* and behavioral testing

Two-month old C57BL6/J mice (18-25 g) of both sexes were injected intraperitoneally (IP) with 20mg/kg paclitaxel (Bristol-Myers Squibb) every other day (days 1, 3, 5, and 7, for a total of 4 injections), while 2 mg/kg 8-Br-cADPR was injected after every two paclitaxel injections (days 4 and 8, for a total of 2 injections). Paclitaxel was prepared in 1 part vehicle (1:1 v/v Cremophor EL [EMD Millipore] and 200- proof ethanol) and 2 parts sterile saline (UPS), and injected at 10μL/g. Control mice were injected with 1 part vehicle and 2 parts saline. 8-Br-cADPR (Sigma) was prepared in sterile saline and injected at 10 µl/g, and control mice were injected with saline. Mice were habituated for two days, and baseline behavioral performance was assessed in the next two days and averaged. The first paclitaxel injection (day 1) was given three days later, and mice were behaviorally tested 9 days after the final injection (day 17). Noxious mechanosensation threshold was assayed as described previously (Pease-Raissi et al., 2017) using Von Frey filaments (0.008-1.4 g). Withdrawal threshold was determined to be the applied force at which the animal withdrew the stimulated paw on at least 2 of 10 applications.

### Tumor-bearing mice

E0771 mouse breast tumor cells (ATCC) were injected (2 x 10^5^ cells/injection) orthotopically into the thoracic fat pads of 2-month old C57BL/6J female mice (Jackson) in 40% matrigel. Seven days after tumor cells injection, paclitaxel or vehicle was injected intraperitoneally (IP) every other day for 16 days (days 1, 3, 5, 7, 9, 11, 13 and 15, for a total of 8 injections; 4 mg/kg for the first two injections and 20 mg/kg for the remaining six injections). 8-Br-cADPR or saline were injected intraperitoneally after every two injections of paclitaxel (days 4, 8, 12 and 16, for a total of 4 injections; 2mg/kg each injection). Tumor size and animal weight were measured before the first paclitaxel injection and every 3-4 days after. Tumor size was assessed by measuring the long and short axes, and volume was calculated by use of the modified ellipsoid formula (long x short^2^ x 0.5). The size of each tumor was normalized to the tumor size acquired on the day before the first injection of paclitaxel.

### Epidermal footpad innervation

Mice hindpaw footpads were prepared as described (Pease-Raissi et al., 2017). Briefly, mice were euthanized with isoflurane and footpad tissue was removed and divided into thick (dermal papillae containing) and thin (non-dermal papillae containing) skin. Footpads were fixed in Zamboni’s fixative overnight at 4°C, cryopreserved in 30% sucrose in PBS for two days at 4°C, frozen, and sectioned into 30 μm floating sections. Sections were permeabilized and blocked with 0.1% Triton X-100 in PBS supplemented with 10% normal goat serum for 1 hour at room temperature and incubated with mouse anti-Tuj1 (1:300; BioLegend) overnight at 4°C. Sections were then incubated with goat anti-mouse AlexaFluor 488 (1:200; Invitrogen) and DAPI (Invitrogen 1:1000) for 2 hours at room temperature and mounted on gelatin-coated slides. IENF images were acquired on a Nikon Ni-E C2 confocal with a 40X 1.3NA oil objective as 30-31μm z-stacks, and converted to maximum intensity projection image for quantification. Intraepidermal nerve fiber density was determined to be the number of Tuj1-positive fibers expanding into the epidermis per 100μm epidermal length.

### Quantification and statistical analysis

Data are expressed as mean +/- SEM. For grouped data multiple comparisons, data were analyzed by two-way ANOVA with Tukey’s multiple comparisons test. For columned data multiple comparisons, data were analyzed by one-way ANOVA with Tukey’s multiple comparisons test. For calcium imaging, data were analyzed by two-way ANOVA and data at each time point were compared using Tukey’s multiple comparisons test. Significance was place at p<0.05. Statistical analysis was done using GraphPad Prism.

## KEY RESOURCES TABLE

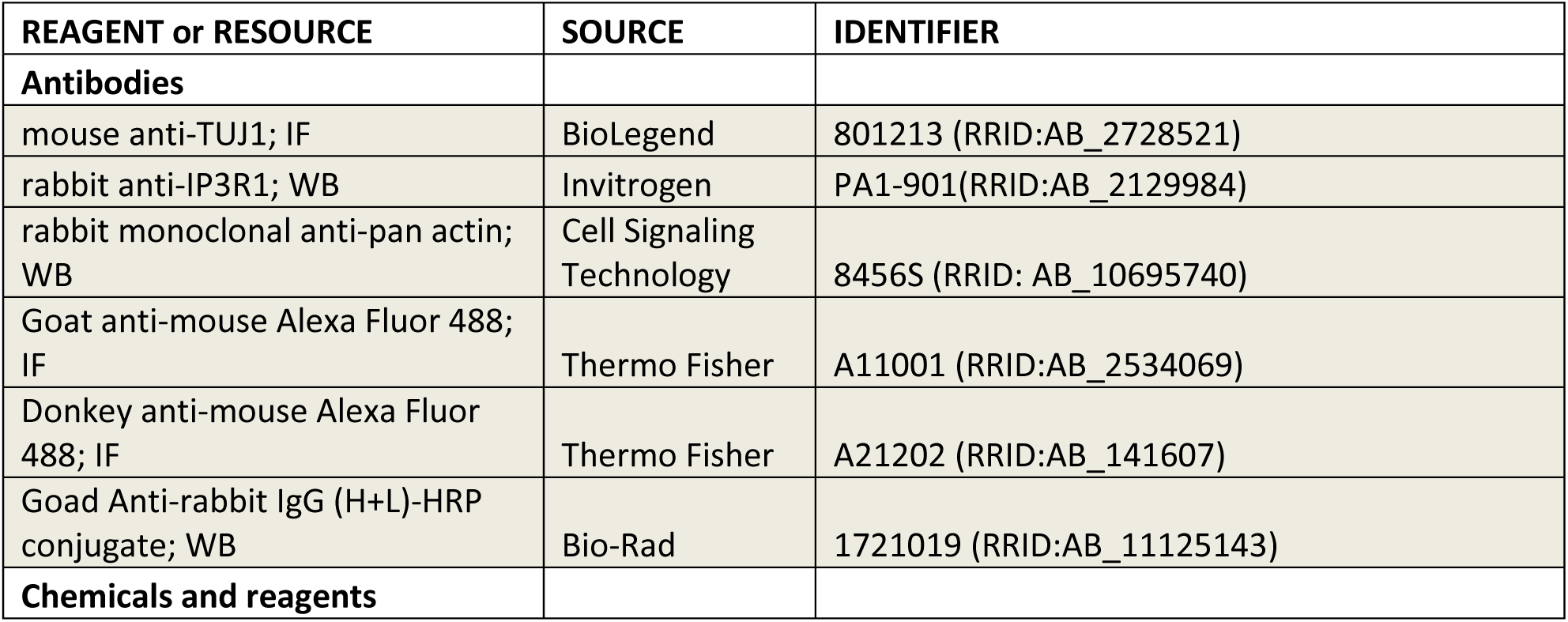
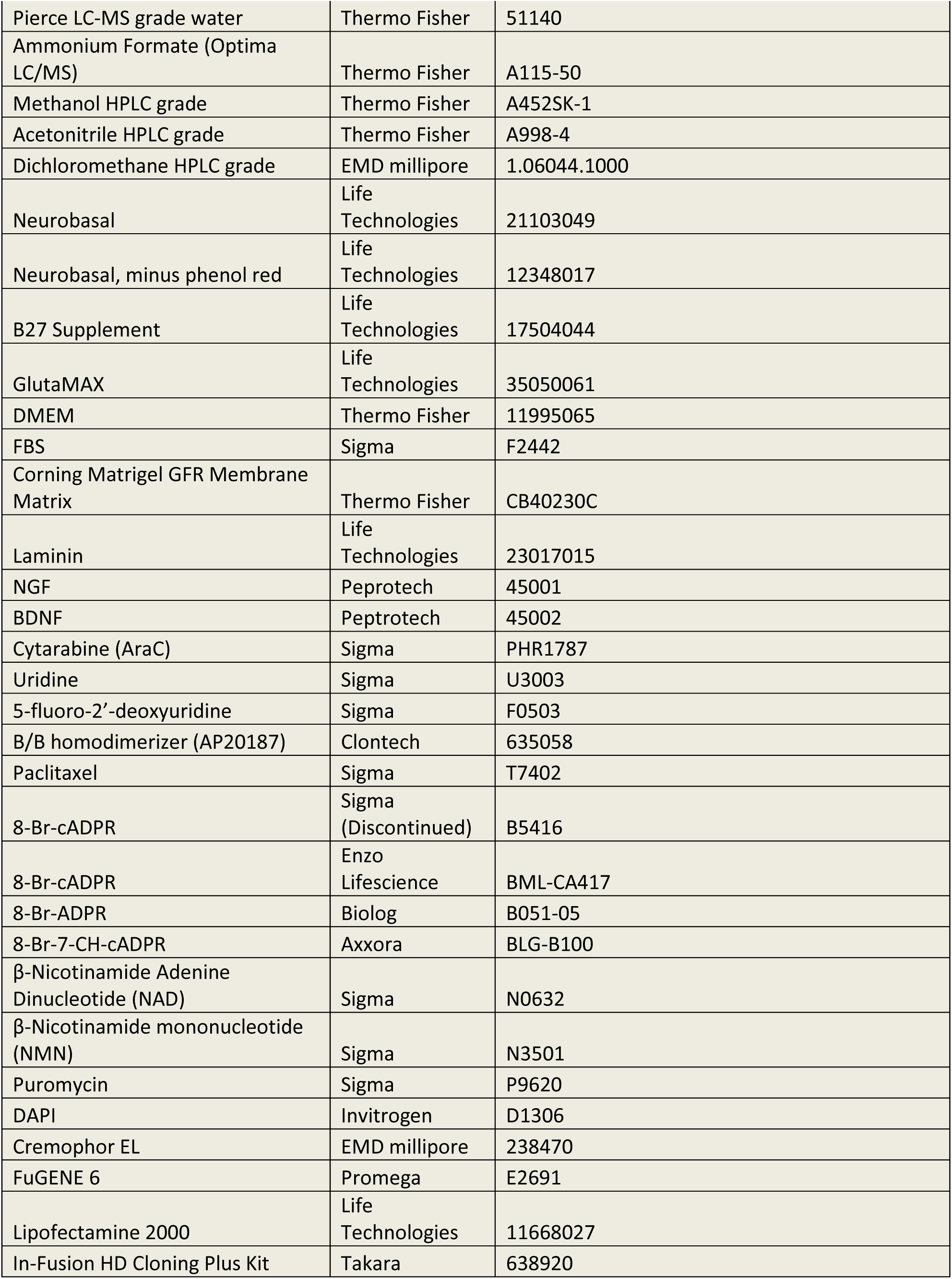
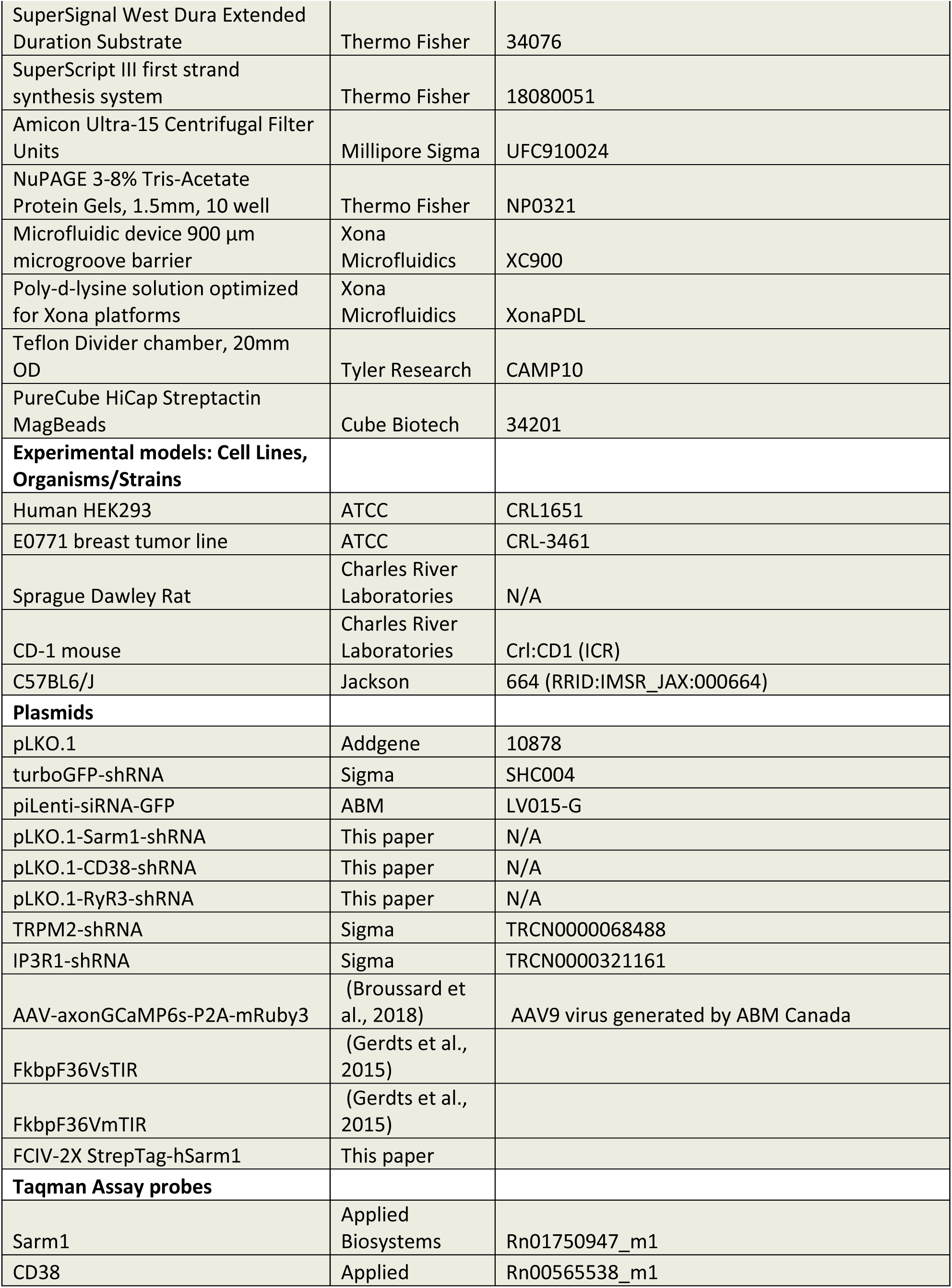
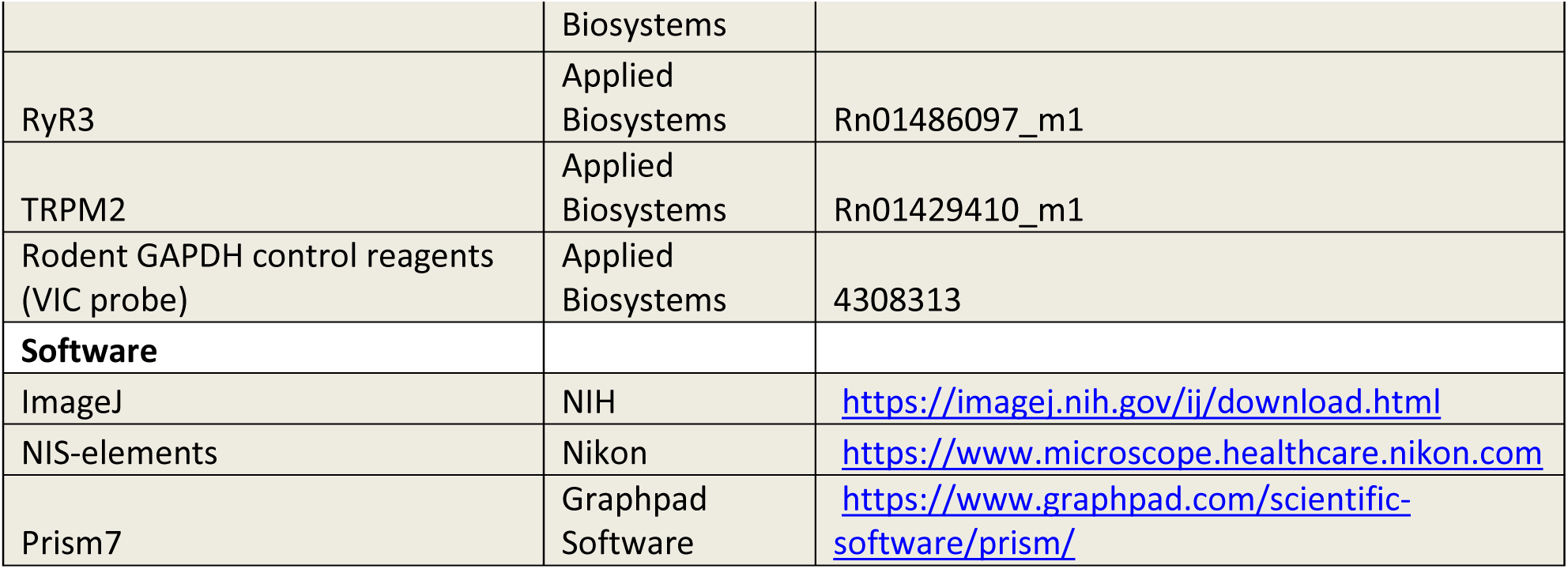

## CONTACT FOR REAGENT AND RESOURCE SHARING

Further information and requests for resources and reagents should be directed to and will be fulfilled by the Lead Contact, Rosalind Segal.

## Author contributions

Y.L., M.F.P.-M., D.A., J.J.Z., A.D., J.M., and R.A.S., designed research.

Y.L., M.F.P.-M., S.T., M.D.S.T.R., D.A., J.S.B., T.J., J.Z., and K.W.K., performed research.

Y.L., M.F.P.-M., S.T., M.D.S.T.R., D.A., J.Z., and K.W.K., analyzed data.

Y.L. and R.A.S. wrote the paper with input from M.F.P.-M., A.D., J.M., J.S.B., J.J.Z., M.D.S.T.R., and D.A..

## CONFLICT OF INTEREST STATEMENT

J.S.B. is a scientific consultant for Geode Therapeutics Inc. T.J. is a scientific consultant for Crimson Biotech Inc. J.J.Z. is a co-founder and board director of Crimson Biotech Inc. and Geode Therapeutics Inc..

A.D. and J.M. are co-founders, scientific advisory board members, and shareholders of Disarm Therapeutics.

## ACKNOWLEDGMENTS

We thank Timothy Walseth (University of Minnesota) for comments and discussion on cADPR functions. We thank Nika Danial, Accalia Fu, M. Carmen Fernandez-Aguera for discussion on metabolic studies. We thank the Segal lab and Stiles lab for helpful comments on the manuscript. This work was supported by the Edward R. and Anne G. Lefler Center Postdoctoral Fellowship to Y.L., Friends of Dana-Farber Cancer Institute grant to J.S.B., Breast Cancer Research Foundation grant to J.J.Z., and the National Institutes of Health grants R35 CA210057 to J.J.Z., R01 CA205255 to R.A.S., R01 CA219866 and RO1 NS087632 to A.D. and J.M

## SUPPLEMENTARY FIGURE LEGENDS

**FigureS1.**
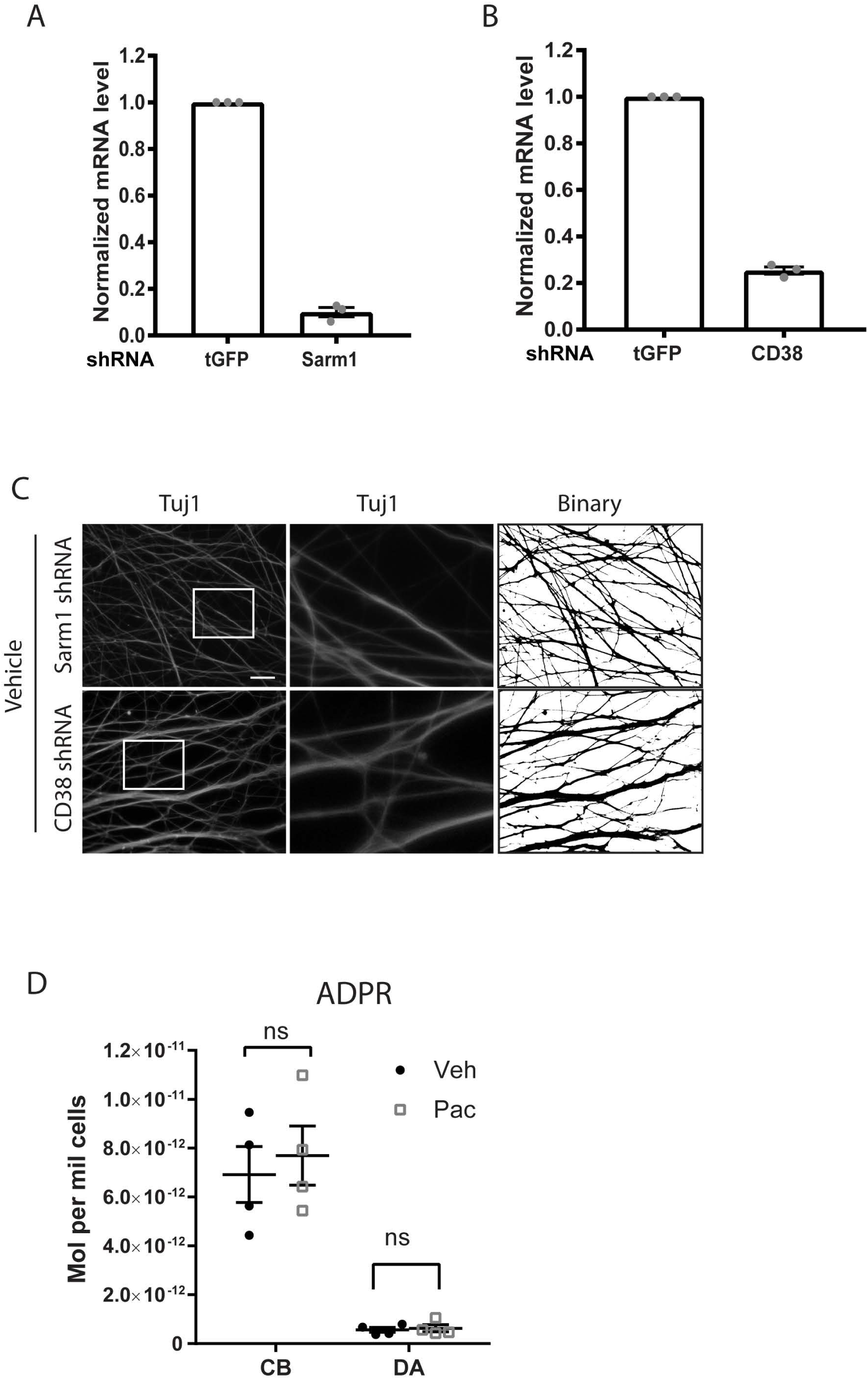
(A-B) Normalized mRNA level of DRG neurons showing knockdown efficiency of Sarm1 shRNA (A) and CD38 shRNA (B) compared to tGFP-control shRNA. Data represents mean +/- SEM; individual data points are shown. N represents three independent experiments. (C) Tuj1 immunostaining and corresponding binarized images of DRG axons transduced with Sarm1 shRNA or CD38 shRNA treated with DMSO (Veh). Scale bar: 20µm. (D) ADPR level in cell bodies (CB) and distal axons (DA) of DRGs after 24hrs of 30nM paclitaxel (Pac) or DMSO (Veh) added to axons. Data represent mean +/- SEM, individual data points are shown. N represents independent experiments.

**FigureS2.**
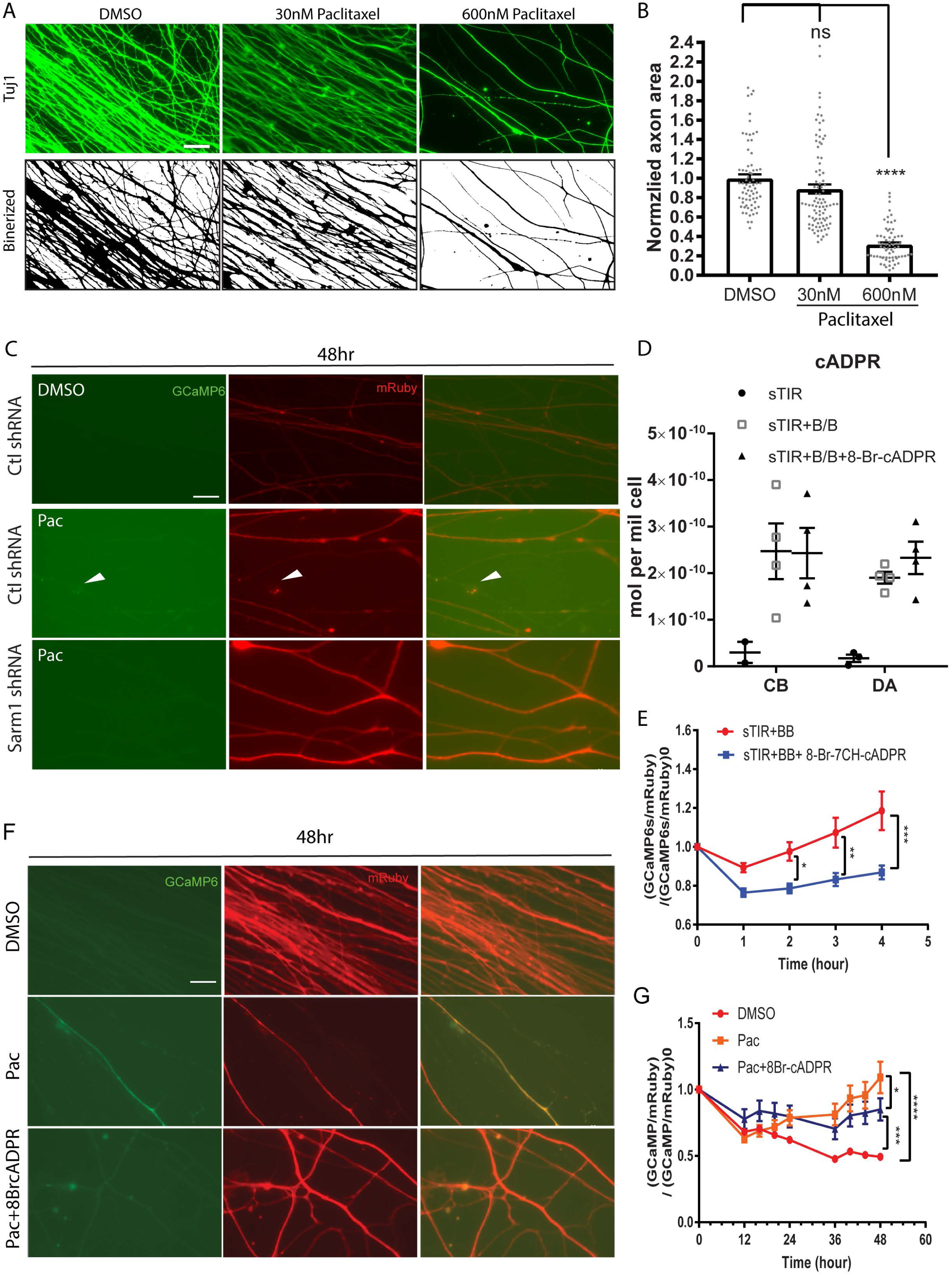
(A) Tuj1 immunostaining and corresponding binarized images of axons of DRG neurons after 48hrs of 30nM or 600nM paclitaxel, or DMSO. Scale bar: 20µm. (B) Normalized area of axons after 48hrs of 30nM, or 600nM paclitaxel or DMSO applied to axons. Data represents mean +/- SEM; individual data points are shown. N represents number of images, with 10-20 images of each sample from 1-2 samples per experiment and 3 independent experiments. ****p<0.0001 by one-way ANOVA with Tukey’s multiple comparison test. (C) Representative images of GCaMP6s (green) and mRuby3 (red) of axons of DRG neurons expressing tGFP shRNA (Ctl) or Sarm1 shRNA after 48hrs of DMSO (Veh) or 600nM paclitaxel (Pac) treatment. White arrow indicates axon with increased calcium at 36hrs, and degenerated at 48hrs (also see Figure2A). Scale bar: 20µm. (D) Level of cADPR in cell bodies (CB) and distal axons (DA) from compartmentalized cultures after 3hrs of 50nM B/B homodimerizer, 50nM B/B homodimerizer together with 10µM 8-Br-cADPR, or vehicle added to axons following infection with lentivirus expressing FkbpF36V-sTIR. Data represent mean +/- SEM; individual data points are shown. For two samples of vehicle treated cell bodies (CB) and one of distal axons (DA), no cADPR was detected. These data points are not shown. N represents four independent experiments. (E) Representative images of GCaMP6s (green) and mRuby3 (red) of DRG axons treated with DMSO, 600nM paclitaxel alone or 600nM paclitaxel with 20µM 8-Br-cADPR. Scale bar: 20µm. (D) Quantification of calcium signal calculated as (GCaMP6s/mRuby3)/(GCaMP6s/mRuby3)_0_. (GCaMP6s/mRuby3) represents calcium signal at each time point, and (GCaMP6s/mRuby3)_0_ represents calcium signal at time 0 before treatment. Data represent mean +/- SEM. N represents number of images. 8-10 images for each sample per experiment. Data pooled from nine independent experiments. *p<0.05, ***p<0.001 by two-way ANOVA with Tukey’s multiple comparison test for each time point.

**FigureS3.**
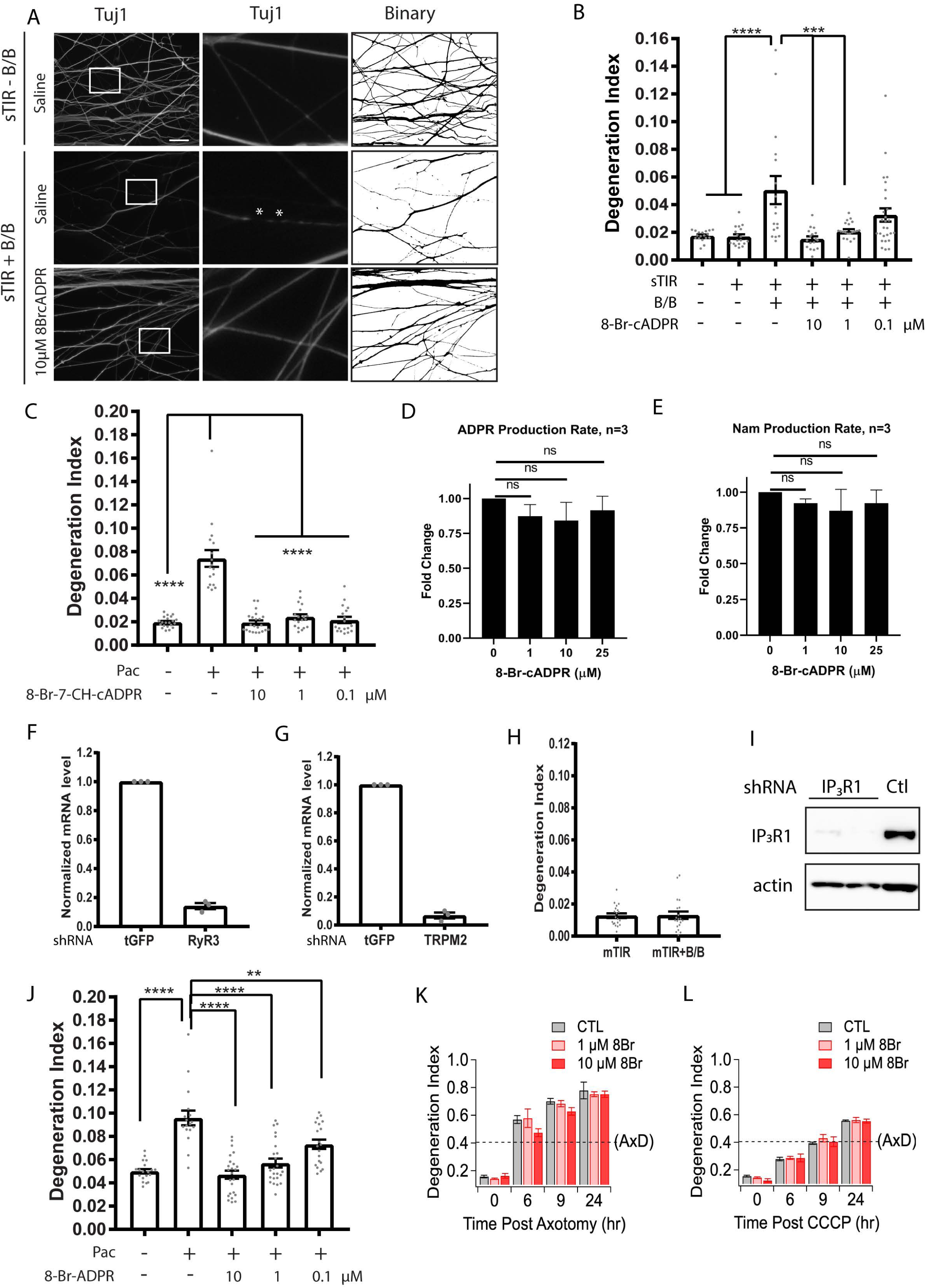
(A) Tuj1 immunostaining and corresponding binarized images of axons of DRGs expressing FkbpF36VsTIR treated with 50nM B/B homodimerizer alone or together with indicated dosage of 8-Br-cADPR. White boxes outline regions shown at higher magnification in the center panels. White stars indicate fragmented region of axons displayed as interruptions in Tuj1 continuity. Scale bar: 20µm. (B) Degeneration index of (A). Data represent mean +/- SEM. Individual data points were shown. N represents number of images. Data pooled from six independent experiments. **p<0.01 ***p<0.001 ****p < 0.0001 by one-way ANOVA with Tukey’s multiple comparison test. (C) Degeneration index of DRG axons after 24hrs of DMSO (Veh) or 30nM paclitaxel (Pac) with indicated concentrations of 8-Br-7CH-cADPR. ****p < 0.0001 by one-way ANOVA with Tukey’s multiple comparison test. Data represent mean +/- SEM. Individual data points are shown. N represents number of images. Data pooled from three independent experiments. (D-E) Production rate of ADPR (D) and nicotinamide (Nam) (E) using purified Sarm1 protein supplemented with NAD and NMN together with indicated dosages of 8-Br-cADPR. Data represents mean +/- SEM. Data pooled from three independent experiments. (F-G) Normalized mRNA levels of DRG neurons showing knockdown efficiency of RyR3 shRNA (E) and TRPM2 shRNA (F). Data represents mean +/- SEM. Individual data points are shown. N represents three independent experiments. (H) Degeneration index of DRG axons transduced with lentivirus expressing FkbpF36V-mTIR with or without 50nM B/B homodimerizer treatment for 3hrs. Data represent mean +/- SEM. N represents number of images. Individual data points are shown. Data pooled from three independent experiments. (I) Western blot probed against IP_3_R1 and pan-actin showing knockdown efficiency of IP_3_R1 shRNA in DRG neurons. (J) Degeneration index of DRG axons after 24hrs of DMSO (Veh) or 30nM paclitaxel (Pac) with indicated concentrations of 8-Br-ADPR. **p<0.01 ****p < 0.0001 by one-way ANOVA with Tukey’s multiple comparison test. Data represent mean +/- SEM. Individual data points are shown. N represents number of images. Data pooled from three independent experiments. (K-L) Degeneration index of DRG axons before and 6, 9 and 24 hours after axotomy (K) or CCCP treatment (L). ****p < 0.0001 by one-way ANOVA with Tukey’s multiple comparison test. Data represent mean +/- SEM. N represents number of images. Data pooled from three independent experiments.

**FigureS4.**
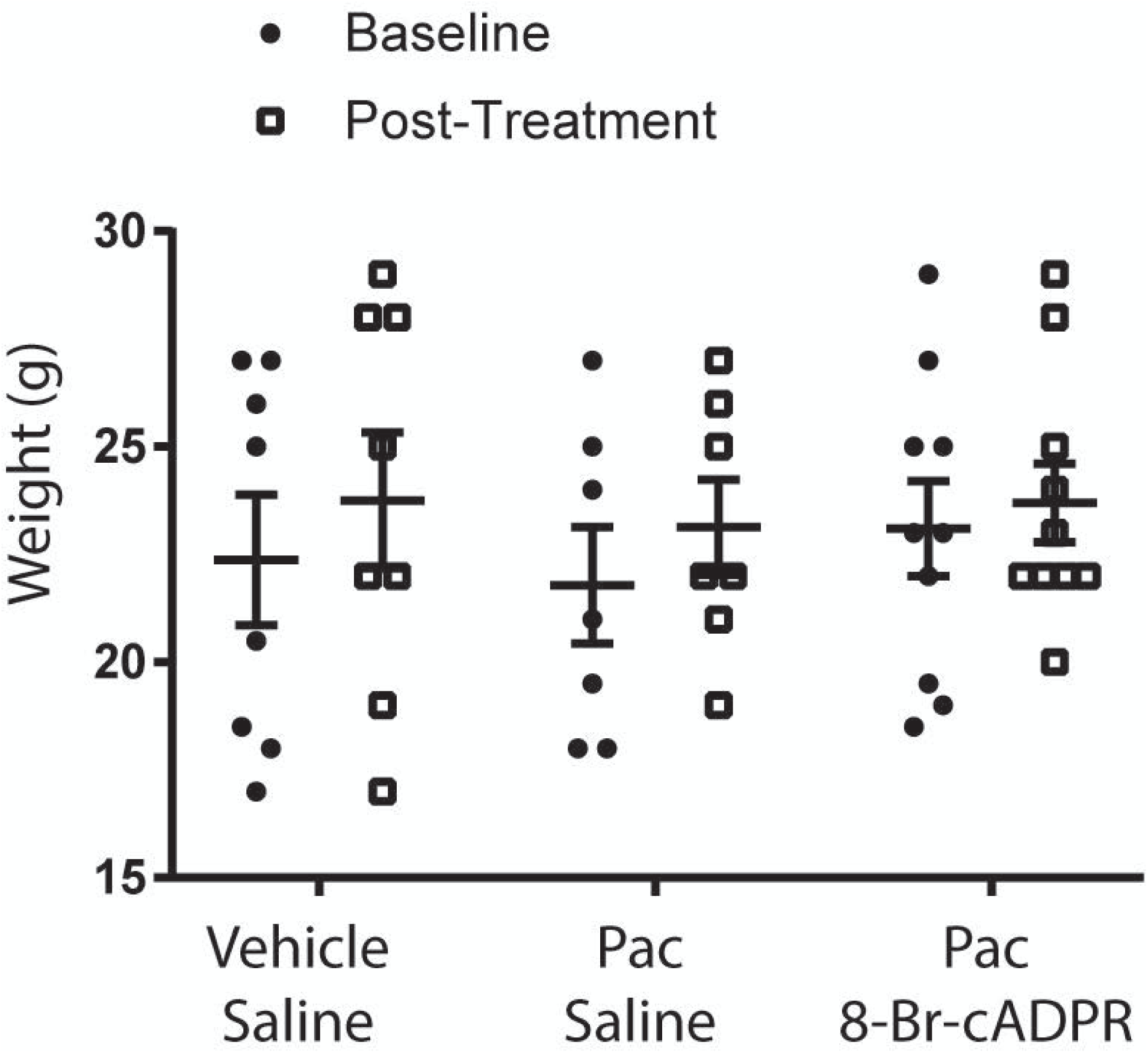
(A) Body weights of animals before (Baseline) and after treatment (Post-treatment). Data represents mean +/- SEM. Individual data points are shown. N represents number of animals.

